# Astrocytic L-lactate signaling in the anterior cingulate cortex is essential for schema memory and neuronal mitochondrial biogenesis

**DOI:** 10.1101/2023.01.08.523156

**Authors:** Mastura Akter, Mahadi Hasan, Aruna Surendran Ramkrishnan, Xianlin Zheng, Zafar Iqbal, Zhongqi Fu, Zhuogui Lei, Anwarul Karim, Ying Li

**Author notes:** **Corresponding author:** Ying Li, MD., Department of Neuroscience, Department of Biomedical Sciences, City University of Hong Kong, Tat Chee Avenue, Kowloon, Hong Kong. **Authors’ email**: Mastura Akter, Mahadi Hasan, Aruna Surendran Ramkrishnan, Xianlin Zheng, Zafar Iqbal, Zhongqi Fu, Zhuogui Lei, Anwarul Karim.

## Abstract

Astrocyte-derived L-lactate was shown to confer beneficial effects on synaptic plasticity and cognitive functions. However, how astrocytic G_i_ signaling in the anterior cingulate cortex (ACC) modulates L-lactate levels and schema memory is not clear. Here, using chemogenetic approach and well-established behavioral paradigm, we demonstrate that astrocytic G_i_ pathway activation in ACC causes significant impairment in flavor-place paired associates (PA) learning, schema formation, and PA memory retrieval in rats. It also impairs new PA learning even if a prior associative schema exists. These impairments were mediated by decreased L-lactate in ACC due to astrocytic G_i_ activation. Concurrent exogenous L-lactate administration bilaterally into the ACC rescues these impairments. Furthermore, we show that the impaired schema memory formation was associated with a decreased neuronal mitochondrial biogenesis caused by decreased L-lactate level in ACC upon G_i_ activation. Our study also reveals that L-lactate mediated mitochondrial biogenesis is dependent on monocarboxylate transporter 2 and NMDA receptor activity – discovering a previously unrecognized signaling role of L-lactate. These findings expand our understanding of the role of astrocytes and L-lactate in brain functions.

## 1. Introduction

Astrocyte, the predominant type of glia in brain, is involved in complex brain functions including learning, memory, and synaptic plasticity (Santello et al., 2019; Doron et al., 2022). They can modulate neuronal activity by releasing and regulating different neuroactive molecules (Ota et al., 2013; Doron et al., 2022). Astrocytes express numerous transporters and receptors including G protein-coupled receptors (GPCRs) to modulate their own as well as neuronal activity. Designer Receptors Exclusively Activated by Designer Drugs (DREADDs) are genetically modified GPCRs which allow researchers to control cellular activity via modulation of GPCR signaling with the application of selective ligands (Urban & Roth, 2015; Yu et al., 2020). These chemogenetic tools have been used to modulate the function of neurons and astrocytes in different brain regions (Armbruster et al., 2007; Alexander et al., 2009; Zhu et al., 2014; Koike et al., 2016; Adamsky et al., 2018; Jones et al., 2018; Durkee et al., 2019; Pati et al., 2019; Kol et al., 2020; Oguchi et al., 2021; Lei et al., 2022; Liu, S. et al., 2022). Using chemogenetic approach, astrocytic G_i_ pathway activation in hippocampus has recently been shown to modulate cognitive functions (Jones et al., 2018; Kol et al., 2020; Liu, S. et al., 2022), although the mechanism is still not fully understood. Moreover, how astrocytic G_i_ signaling in the anterior cingulate cortex (ACC) affects cognitive functions – particularly schema memory - is yet unknown.

L-lactate is a metabolic end product of glycolysis and works as an energy substrate for various tissues including the brain. According to the Astrocyte-Neuron L-lactate Shuttle (ANLS) hypothesis (Pellerin & Magistretti, 1994; Magistretti & Allaman, 2018), L-lactate is produced by astrocytes through glycogenolysis and glycolysis and then transported into the neurons through monocarboxylate transporter 2 (MCT2) to fuel the high metabolic demand in neurons to maintain various physiological activities including neural plasticity and memory formation (Magistretti & Allaman, 2018). However, a recent study has argued that the energy demand during neuronal activation is fueled by the glucose rather than astrocytes-derived L-lactate (Diaz-Garcia et al., 2017). Nevertheless, multiple studies clearly demonstrated that L-lactate confers beneficial effect in learning and memory (Newman et al., 2011; Suzuki, Akinobu et al., 2011; Wang, J. et al., 2017; Harris et al., 2019; Netzahualcoyotzi & Pellerin, 2020; Vezzoli et al., 2020). Administration of L-lactate into the hippocampus enhanced memory in rats whereas inhibition of astrocytic glycogenolysis or inhibition of astrocytic or neuronal MCTs in the hippocampus impaired memory formation (Newman et al., 2011; Suzuki, A. et al., 2011; Netzahualcoyotzi & Pellerin, 2020). However, how L-lactate in ACC affects schema memory is yet unknown.

Using a behavioral paradigm, Tse *et al*. showed that learning of multiple flavor-place paired associates (PAs) leads to the development of cortical associative schema in rats that allows rapid assimilation of new PAs into the existing schema (Tse et al., 2007; Tse et al., 2011). Previously, our team showed that bilateral infusion of lidocaine (a neuronal blocker) into hippocampus or ACC prevents PA learning, schema formation, and memory retrieval (Hasan et al., 2019). The study also showed increased oligodendrogenesis and adaptive myelination in the ACC of rats after repeated PA training. Furthermore, it demonstrated that myelination in ACC is necessary for PA learning and memory retrieval suggesting important role of oligodendrocytes in schema memory. However, role of astrocytes of ACC in PA learning, schema formation, and memory retrieval is still unknown. Using hM4Di (a G_i_-coupled GPCR) DREADD, here we show that astrocytic G_i_ pathway activation in ACC causes significant impairment in PAs learning, schema formation, and memory retrieval in rats. We also show that the impairment is mediated by a decrease in L-lactate level in the ACC upon astrocytic G_i_ activation. Concurrent exogenous L-lactate administration into ACC rescues these impairments. Furthermore, we discover that astrocytic G_i_ activation diminishes neuronal mitochondrial biogenesis which could be rescued by exogenous L-lactate and that L-lactate induced neuronal mitochondrial biogenesis requires MCT2 and NMDAR activity – revealing a previously unrecognized L-lactate signaling mechanism in controlling neuronal mitochondrial biogenesis.

## 2. Methods

### 2.1. Animal use and care

Adult male Sprague-Dawley rats weighting about 250-300 g were used in this study. All rats were housed in a standard laboratory facility (25°C, 50% humidity, 12-hours light/dark cycle with light on at 7.00 AM). All animals were supplied by the Laboratory Animal Services Centre, Chinese University of Hong Kong. All experimental procedures were conducted according to the guidelines developed by the National Institute of Health (NIH, Bethesda, MD, USA) and were approved by the Committee on Use and Care of Animals at City University of Hong Kong and the licensing authority from the Department of Health of Hong Kong [Reference no. (22-2) in DH/HT & A/2/5 Pt.8] to conduct experiments. Rats were provided with food and water *ad libitum* except for the period of schema experiments when food restriction was applied.

### 2.2. Paired-associate behavioral protocol

#### 2.2.1. Paired-associate experiment design

We used a behavioral paradigm of multiple flavor-place paired-associates (PAs) learning as described previously (Tse et al., 2007). The event arena, as shown in Supplementary Fig. S1A, contains four start boxes and multiple sand wells. A flavored food pellet (flavor cue) is given in the start box, and a specific sand well (place cue) contains three more of that flavored food pellet at the bottom of it. There are multiple specific flavor-place PAs, for example beef flavor is paired with sand well number 1, strawberry flavor is paired with sand well number 2, and so on. When a rat is placed in a start box that contains a specific flavor cue, they need to use spatial memory to find out the correct sand well that contains that specific flavored food.

Our experimental set up and timeline is illustrated in Supplementary Fig. S1B. We habituated rats for five days (session −7 to −3) so that they become familiar with the event arena and learn digging sand wells. Then, we conducted pretraining for two days (session −2 and −1) to introduce them to the six original flavor-place PAs (OPAs). After that, session 1-18 were conducted as 4-5 sessions/week. In each of the sessions (S) of 1, 2, 4-8, 10-17, each rat was trained with six PAs, and the normal control rats are expected to learn the flavor-place associations so that if a flavor cue is given in the start box, they should be able to find out the correct cued sand well to get more of that flavored food. S3, S9, S18 were non-rewarded probe tests (PT) where rats were given a flavor cue at the start box, but the cued sand well did not contain any food pellet. After getting the flavor cue at the start box, rats were given 120 seconds to find out the cued sand well. PTs reflect memory retrieval. If a rat can retrieve PAs memory well, it will spend more time in digging the cued sand well. In S19, two new PAs (NPA7 and 8) were introduced by replacing two old PAs (PA1 and PA6 respectively). The normal control rats, using the existing schema developed from previous sessions, are expected to learn the new PAs in this single session. This session was followed by S20 which is a non-rewarded PT, where the learning and memory retrieval of the new PAs learned from S19 were tested. If a rat learns the new PAs introduced in S19 well, it will spend more time in digging the new cued sand well.

#### 2.2.2. Performance measures in PA training sessions

*Performance index (PI):* It was calculated for each rat with the following formula:

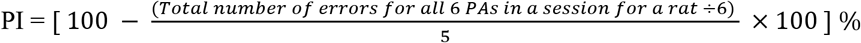

Errors is the number of incorrect (non-cued) sand well(s) the rat dug before digging the correct well (cued). Digging was defined as displacement of sand around sand well by rat.

#### 2.2.3. Performance measures in PT1 (S3), PT2 (S9), PT3 (S18), and PT7 (S34)

Digging time (out of 120 seconds) in the cued and non-cued sand wells were measured. Then proportion of time spent in digging the cued and non-cued wells in respect to the total digging time was calculated as follows:

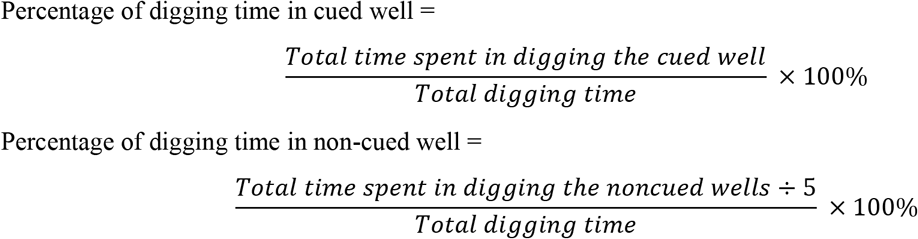

#### 2.2.4. Performance measures in PT4 (S20), PT5 (S23), PT6 (S28)

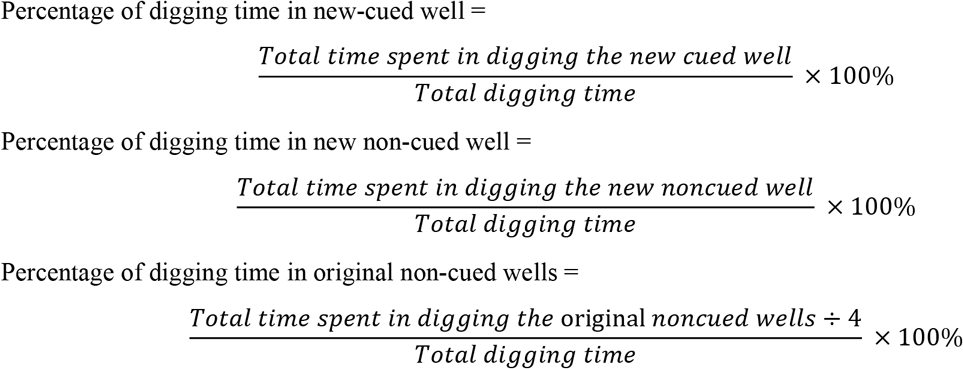

### 2.3. Stereotactic surgical procedures, viral vector injection, and CNO administration

To express hM4Di in the ACC astrocytes, AAV8-GFAP-hM4Di-mCherry was used (original viral titer 3×10^12^ vg/ml diluted in 1:10 in PBS, Obio Technology (Shanghai) Corp. Ltd). Rats were anesthetized with 50 mg/kg sodium pentobarbital (Dorminal 20%, Alfasan International BV, Woerden, Holland) administered intraperitoneally (I.P.) and placed in a stereotaxic frame. After exposing the skull, bilateral craniotomy was done (0.5 - 0.8 mm holes, 2.4 mm anterior to bregma, 0.5 mm lateral from midline). A 10 μl micro-syringe (Hamilton, NV, USA) with a 33-gauge metal needle was used to perform the microinjections. We injected 400 nl of viral vector bilaterally into the ACC regions (2-3 mm ventral from the surface of the skull at the craniotomy site) with injection flow rate of 0.1 μl/min (controlled by microinjection pump, World Precision Instruments, USA). The needle was left in place for an additional five mins after the injection was completed. Then it was slowly withdrawn. After withdrawing the needle, the scalp was sutured, and immediate postoperative care was provided with local anesthetic (xylocaine, 2%) applied to the incision site for analgesia and allowing the rats to recover from anesthesia under a heat pad. The rats were returned to their home-cage after awaking. All rats were allowed three weeks of rest to ensure hM4Di expression.

Clozapine-N-oxide (CNO) dihydrochloride (Hello Bio, Avonmouth, UK, cat. HB6149), a synthetic ligand to activate hM4Di was dissolved in 0.9% NaCl and was injected I.P. at a dose of 3 mg/kg body weight. This dose did not produce any seizure in rats.

### 2.4. Chronic ACC Cannulation for drug infusion

Rats were anesthetized with 50 mg/kg I.P. sodium pentobarbital administration. Stainless steel guide cannulae (Double/O.D.0.41mm-27G/C, cat # 62069, RWD life science.com.ltd.) were bilaterally positioned into ACC region (2.4 mm anterior to bregma and 0.5 mm lateral from midline, 2 mm dorso-ventral from skull surface). The guide cannulae were fixed to the skull with dental cement (mega PRESSNV + JET X, megadental GmbH, Budingen, Germany). Dummy cannulae (Cat # 62169, RWD life science.com.ltd.) 0.5 mm longer than the guide cannulae, were inserted into the guide cannulae to prevent blockage and reduce the risk of infection. The rats were provided a minimum recovery period of one week before the other experimental procedures.

Drugs were administered bilaterally into ACC at a flow rate of 0.333 μl/min using a 33-gauge internal injecting needle. Drugs and their doses per ACC: 10 nmol L-lactate (Sigma Aldrich, Cat #: L7022) in 1 μl ACSF (Wang, J. et al., 2017); 0.5 μl of 30 mM D-(-)-2-Amino-5-Phosphonopentanoic acid (D-APV, Sigma Aldrich, Cat #: A8054) dissolved in ACSF (Wang, S. H. et al., 2012); 1 μl of 0.5 nmol α-Cyano-4-hydroxycinnamic acid (4-CIN, Sigma Aldrich, Cat #: C2020) dissolved in ACSF containing DMSO (Wang, J. et al., 2017). D-APV and 4-CIN were injected 15 mins before L-lactate infusion where appropriate. The needle was kept in place for additional 5 minutes to allow proper diffusion of L-lactate.

### 2.5. Measurement of cAMP and L-lactate levels

To investigate the effect of ACC astrocytic G_i_ pathway activation on cAMP and L-lactate levels in the ACC, 16 rats were habituated and pre-trained for PA experiment as shown in the Fig. 3A. Then bilateral AAV8-GFAP-hM4Di-mCherry injection into the ACC was done as described before in all rats. In addition, a micro-dialysis guide cannula (CMA Inc.) was inserted into the right sided ACC (2.5 mm ventral from the surface of the skull at the craniotomy site) in the rats that was used for microdialysis later (eight rats). After three weeks, all rats were trained for two sessions with six PAs. For microdialysis in the next session (S3), rats were given I.P. CNO (3 mg/kg body weight) (n=4 rats) or saline (n=4 rats) and placed in the PA even arena. ECF from ACC was collected before, 20, 40, and 60 mins after CNO or saline administration. For collecting ECF, a microdialysis probe which is a Y-shaped catheter containing an inlet and outlet port with a fibrous, semi-permeable membrane at the bottom tip (CMA 11 elite, 3 mm membrane) was inserted into guide cannula. One Fluorinated Ethylene Propylene (FEP) tube (ID 0.12 mm, CMA inc,) was connected to the inlet port and another FEP tube was connected to outlet port. Through inlet FEP tube, artificial cerebrospinal fluid (ACSF, Harvard Apparatus, cat #: 597316) was infused into ACC to maintain artificial neurotransmitter concentration gradient. Through the outlet tube, extracellular fluid from ACC was collected by micro-infusion pump (WPI). For IHC staining of cAMP in S3, rats were given I.P. CNO (3 mg/kg body weight) (n=4 rats) or saline (n=4 rats). After 30 mins, PA training was started, and the rats were sacrificed at 60 mins of CNO/saline administration.

**Fig. 1.**
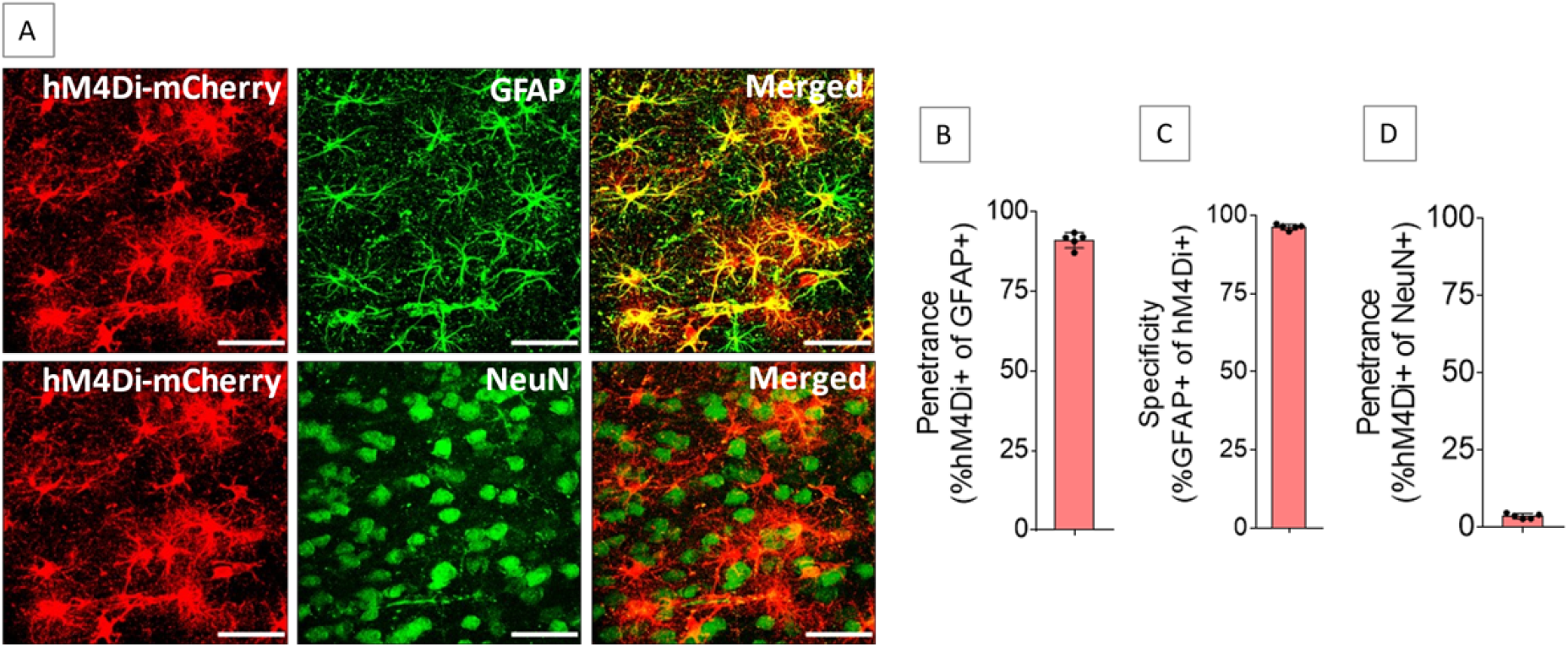
Expression of hM4Di in ACC astrocytes. Injection of AAV8-GFAP-hM4Di-mCherry into ACC resulted in expression of hM4Di (**A**) in 91.1 ± 2.4% of GFAP-positive cells (**B**) with 96.3 ± 0.9% specificity (**C**), whereas 3.6 ± 0.8% of NeuN-positive cells expressed hM4Di (**D**). n=5 rats. Scale bars: 50 μm.

**Fig. 2.**
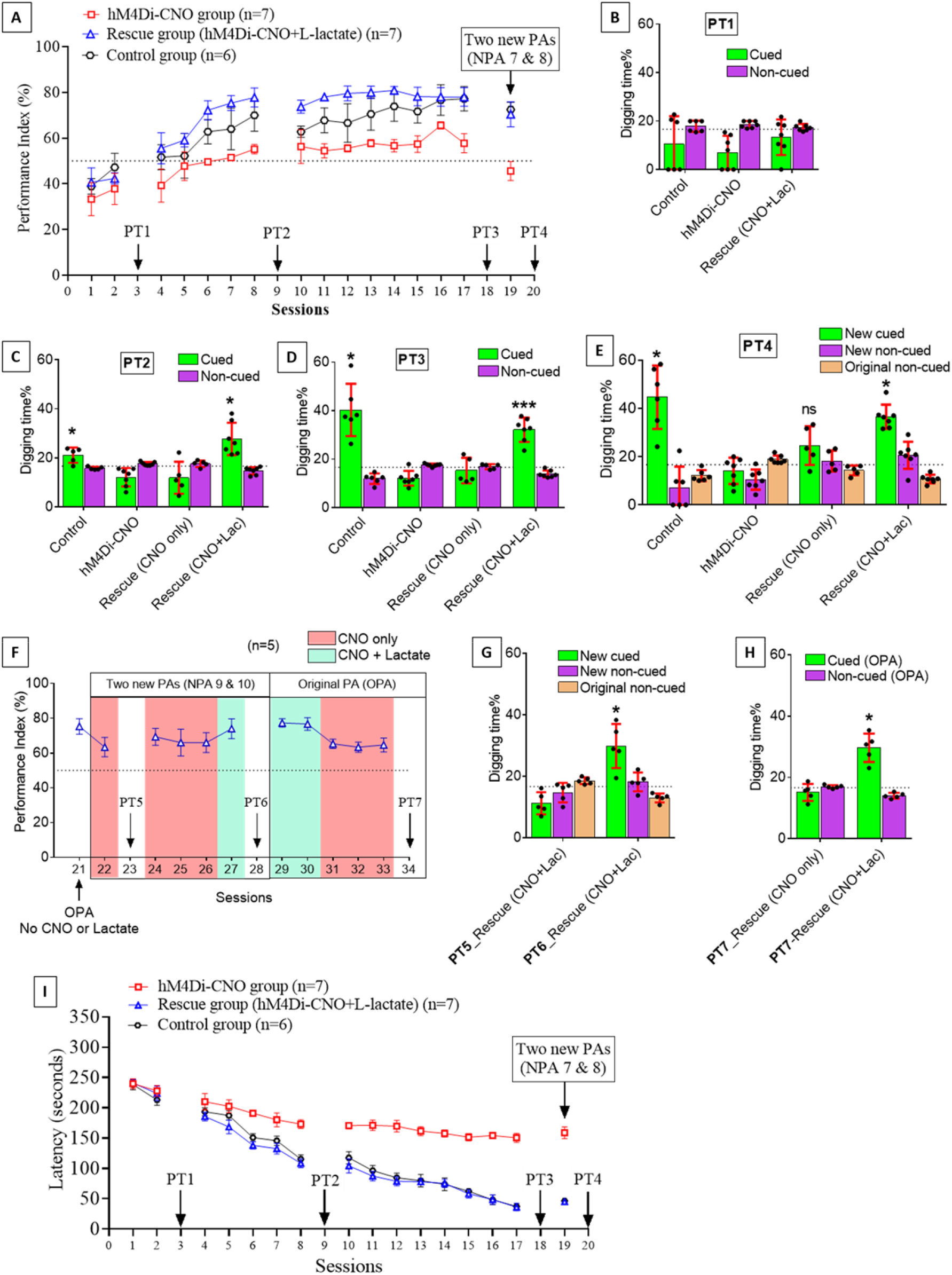
Astrocytic G_i_ pathway activation in ACC impairs PA learning, schema consolidation, memory retrieval, and assimilation of new PAs into existing schema whereas L-lactate can rescue these impairments. **A.**PI (mean ± SD) during the acquisition of the original six PAs (OPAs) (S1-2, 4-8, 10-17) and new PAs (NPAs) (S19) of the control (n=6), control-CNO (n=4), hM4Di-CNO (n=7), and rescue (hM4Di-CNO+L-lactate) (n=7) groups. From S6 onwards, hM4Di-CNO group consistently showed lower PI compared to control. However, concurrent L-lactate administration into ACC (rescue group) can rescue this impairment. Data of control-CNO group shows that CNO application itself has no effect on PA learning. **B, C, & D.** Non-rewarded PTs (PT1, PT2, and PT3 done on S3, S9, and S18, respectively) to test memory retrieval of OPAs for the control, hM4Di-CNO, and rescue groups. The percentage of digging time at the cued location relative to that at the non-cued locations are shown (mean ± SD). Both PT2 and PT3 were done twice for the rescue group. One PT was done in the morning with I.P. CNO administration 30 mins before the PT (CNO only). The other PT was done in the afternoon with I.P. CNO 30 mins before and bilateral L-lactate administration 15 mins before the PT (CNO+Lac). In both PT2 and PT3, the control group spent significantly higher time in digging the cued sand well above chance level indicating that the rats had learned OPAs and could retrieve it. Contrasting to this, hM4Di-CNO group did not spend higher time in digging the cued sand well above chance level. The rescue group showed results similar to hM4Di-CNO group if CNO is given without L-lactate. On the other hand, they showed results similar to the control group if L-lactate is concurrently given with CNO indicating that this group had learned OPAs and could retrieve it. ns=not significant, \**p* < 0.05, and \*\*\**p* < 0.001, one sample t-test comparing the proportion of digging time at the cued sand well with the chance level of 16.67%. **E.** Non-rewarded PT4 (S20) which was done after replacing two OPAs with two NPAs (NPA 7 & 8) in S19 for the control, hM4Di-CNO, and rescue groups. Results show that the control group spent significantly higher time in digging the new-cued sand well above chance level indicating that the rats had learned the NPAs from S19 and could retrieve it in this PT. Contrasting to this, hM4Di-CNO group did not spend higher time in digging the new-cued sand well above chance level. The rescue group showed results similar to hM4Di-CNO group if CNO is given without L-lactate. On the other hand, they showed results similar to the control group if L-lactate is concurrently given with CNO indicating that this group had learned NPAs from S19 and could retrieve it. ns=not significant, and \**p* < 0.05, one sample t-test comparing the proportion of digging time at the cued sand well with the chance level of 16.67%. **F, G, and H.** Continuation study (S21-S34) with the rescue group (n=5). The PI (mean ± SD) is shown in **F**. PT5 and PT6 (done at S23 and S28, respectively) are shown in **G**. PT7, which was done twice, is shown in **H**. In S21, PI was 75.3% for the six OPAs without CNO or L-lactate. For S22-S28, two OPAs was replaced with two NPAs (NPA 9 and 10). In S22, which was done with CNO only, PI dropped to 63.3%. PT5 (**G**) confirms that the rats did not learn the NPA 9 and 10 from S21. In S24-S26, which were done with CNO only, PI remained similarly low (69.3%, 66%, and 66%, respectively) indicating that the rats were not learning the NPAs 9 and 10 despite multiple sessions. In S27, which was done with CNO+L-lactate, PI raised to 77.3% suggesting that they have learned the NPAs in this session. This was confirmed by PT6 (**G**) which showed that they spent significantly higher time in digging the new-cued sand well above chance level. In S29-S34, the six OPAs were restored. Studies in these sessions showed that PI drops from ~77% to ~64% even for the OPAs if L-lactate is not given concurrently with CNO. Furthermore, PT7 (S34) (**H**) shows that astrocytic G_i_ pathway activation in ACC impairs memory retrieval of existing associative schema which can be rescued by administering L-lactate concurrently. ns=not significant, and \**p* < 0.05, \*\**p* < 0.01 and \*\*\**p* < 0.001, one sample t-test comparing the proportion of digging time at the cued sand well with the chance level of 16.67%. **I.** Latency (in seconds) before commencing digging at the correct well for all groups. Data shown as mean ± SD.

**Fig. 3.**
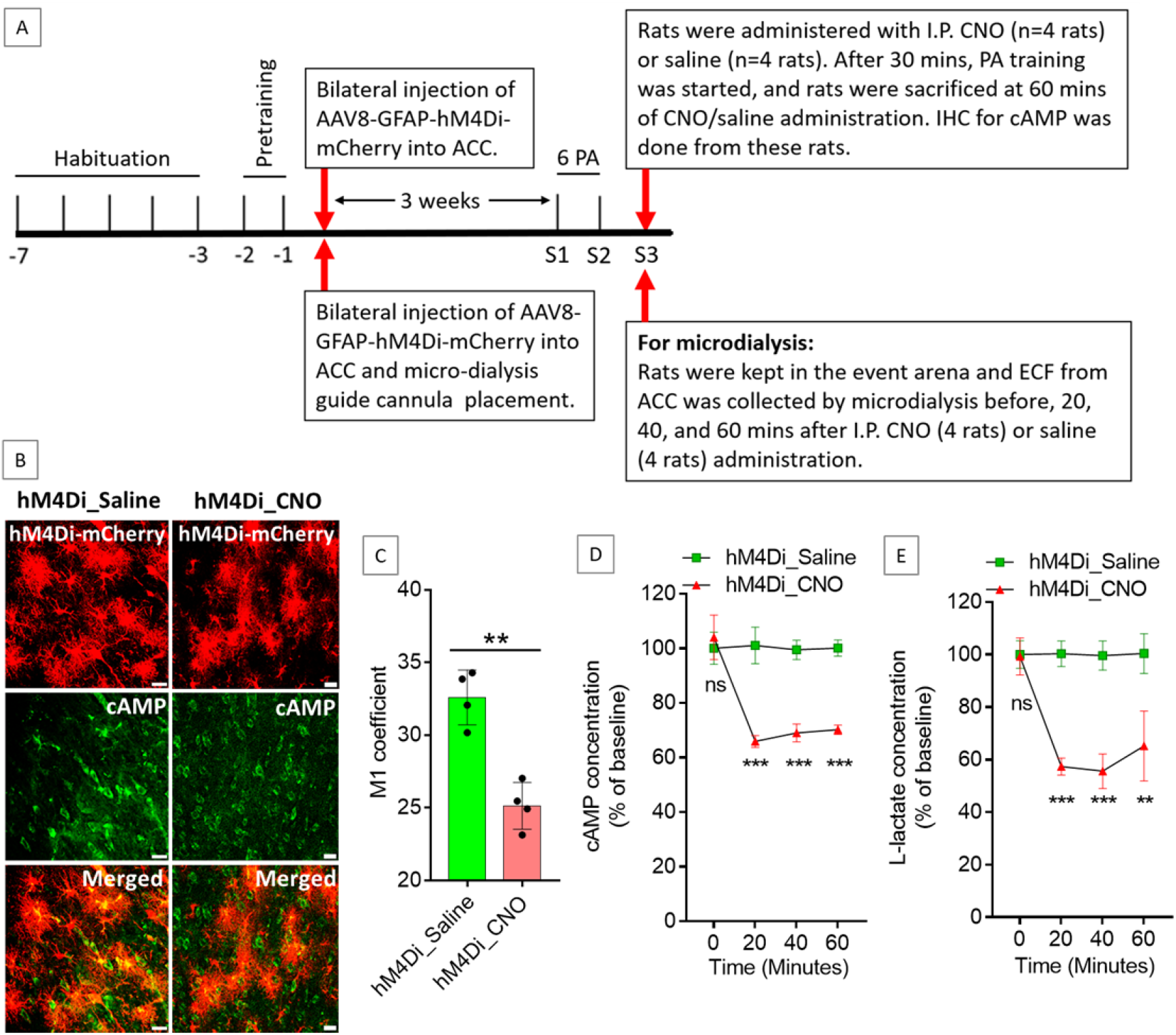
Effect of ACC astrocytic G_i_ activation on cAMP and L-lactate levels in ACC. **A.** Experimental design to investigate the effect of G_i_ activation of ACC astrocytes on cAMP and L-lactate levels. **B, C.** CNO decreases cAMP in the hM4Di expressed cells. (**B**) Confocal micrograph of ACC 60 minutes after intraperitoneal administration of saline or CNO in hM4Di expressed rats. Scale bars: 20 μm. (**C**) Colocalization analysis showing decreased Mander’s coefficient M1 (ratio of cAMP intensity colocalized with hM4Di-mCherry to total cAMP intensity) in CNO administered rats (n=4 in each group). \*\**p* = 0.001, unpaired Student’s t-test, t=6.01, df=6. **D, E.** Microdialysis measurement of cAMP (**D**) and L-lactate (**E**) levels in the ECF of ACC before, 20 mins, 40 mins, and 60 mins after intraperitoneal saline or CNO administration in hM4Di expressed rats (n=4 in each group). \*\*\**p* < 0.001, \*\**p* < 0.01, ns=not significant, unpaired Student’s t-test.

The dialysate collected from ACC were kept at −80 °C until further use. cAMP complete ELISA kit (Abcam, USA, cat. # ab133051) was used to determine the cAMP concentration in ACC dialysate according to the manufacturer’s protocol. Lactate Fluorescence Assay kit (Abcam, USA, cat. # ab65331) was used to determine the L-lactate concentration from the same ACC dialysate according to the manufacturer’s protocol.

### 2.6. Immunohistochemistry and confocal microscopy

After completing experiments, rats were anesthetized by urethane (1.5 g/kg, I.P.) and perfused transcardially with ice-cold PBS for approximately five minutes and then perfused with 4% paraformaldehyde (PFA). The whole brain was taken out and postfixed in 4% PFA overnight at 4°C and cryoprotected in 30% sucrose dissolved in 1X PBS for an additional 3 days at 4°C. The brains were then stored in OCT medium at - 80°C until further use. For IHC, each brain was sectioned at 40 μm using cryostat (Leica, USA) and processed as free-floating sections. Six to eight sections were selected for staining per rat. Sections were incubated with blocking solution of Triton X-100 (0.3% [v/v]) and 10% normal goat serum (NGS) in 0.01 M PBS for 1 hour at room temperature after a brief wash. Then sections were incubated with primary antibodies (listed in Supplementary Table-1) in blocking solution for overnight at 4°C. In the following day, slices were washed 3 times (5 minutes each) and incubated with targeted Alexa flour secondary antibodies (1: 300) in DAPI (4’,6-diamidino-2-phenylindole) for 2 hours at room temperature. Then the sections were mounted into microscopic slides (Epredia^™^ SuperFrost Plus^™^ Adhesion Microscopic Slides) and covered with coverslips (Eprdia Cover Slip) along with fluorescent mounting medium (DAKO). The imaging was done by inverted laser scanning confocal microscope (LSM 880; Carl Zeiss, Oberkochen, Germany). The confocal images for quantitative analysis were acquired under 20X or 40X oil-immersion objectives. Z-stack confocal images of at least 3-6 sections from each rat were obtained.

### 2.7. Relative mitochondrial DNA content quantification

After completing experiments, rats were anesthetized by urethane (1.5 g/kg, I.P.) followed by decapitation. Brain was sectioned on an anodized aluminum brain slicer (Braintree Scientific, Braintree, MA, USA) and ACC were dissected from the sections and kept at −80°C. Total genomic DNA was extracted from ACC using QIAamp DNA Mini Kits (catalogue number 51304) according to manufacturer’s protocol. Genomic DNA quality and quantity were checked with NanoDrop^™^ 1000 Spectrophotometer (Thermo scientific). The DNA sample was stored at −20°C until further use. Quantitative real-time PCR was performed with the SsoAdvanced Universal SYBR Green Supermix (cat# 1725270) using Applied Biosystems QuantStudio™ 3 Real-Time PCR Systems. β-actin gene and mitochondrial D-loop were used as nuclear DNA (nDNA) and mtDNA, respectively, to investigate the abundance of mtDNA relative to nDNA. The primer sequences (Branda et al., 2002; Chou et al., 2007) and reaction mixture protocol is given in Supplementary Table-2. Thermal cycling was done according to the SsoAdvanced Universal SYBR Green Supermix protocol. DNA from each rat was amplified as triplicate. After obtaining both mtDNA and nDNA Ct values from Real Time PCR software, Ct values were averaged from triplicates of each rat. To determine the mtDNA content relative to the nDNA, the following equation (Rooney et al., 2015) was used:

Relative mitochondrial DNA content = 2 × 2^(nDNA Ct – mtDNA Ct)^

### 2.8. Western-blot analysis

The samples were homogenized in mixture (100:1) of RIPA buffer (150 mM NaCl, 50 mM Tris pH 7.4, 1% Triton X-100, 0.1% SDS, 1% sodium deoxycholate) and phosphatase and protease inhibitor cocktail (Sigma-Aldrich). Homogenates were then centrifuged at 14000g for 30 minutes at 4°C. The supernatants were collected carefully as total protein and then Bradford assay was used to determine the protein concentration. Using sodium dodecyl sulphate-polyacrylamide gel electrophoresis (SDS-PAGE), 20 μg of proteins were separated, and then transferred to a polyvinylidene fluoride membrane (PVDF). Then the membrane was incubated with 5% non-fat milk in TBST (containing 0.1% tween 20) for 1 hour followed by incubation with primary antibodies with reference concentration (Supplementary Table-1) for overnight at 4°C. Next day, membranes were washed for 3 times (5 minutes each) with TBST and incubated for 2 hours with horseradish peroxidase coupled secondary goat anti-rabbit or anti-mouse IgG (1:5000, Invitrogen) in TBST. Then membranes were washed 3 times (5 minutes each). Western Bright ECL HRP substrate (Advansta, Inc, cat# K12045-D20) was used to visualize the blot and quantified by FIJI ImageJ software (National Institutes of Health, Bethesda, MD, USA). Expression of target proteins were normalized with that of β-actin level.

### 2.9. Data analysis

Data analyses were done with Prism v.7.0 (GraphPad Software, La Jolla, CA, USA) or MS Excel. Data are presented as mean ± SD as appropriate. Comparisons of continuous data were done with two-tail Student’s t test where appropriate. Image analysis was done with ImageJ. Figures were generated with Prism v.7.0 and Inkscape.

## 3. Results

### 3.1. Expression of hM4Di in ACC astrocytes

ACC of both sides were injected with Adeno-Associated Virus serotype 8 (AAV8) vector encoding mCherry-tagged hM4Di under the control of Glial Fibrillary Acidic Protein (GFAP) promoter to drive hM4Di expression in ACC astrocytes (AAV8-GFAP-hM4Di-mCherry). The hM4Di is a modified human muscarinic receptor M4 that has been engineered to be insensitive to the endogenous ligand acetylcholine but can be activated by its selective ligand clozapine-N-oxide (CNO) (Armbruster et al., 2007). Injection of AAV8-GFAP-hM4Di-mCherry into ACC resulted in expression of hM4Di in ACC astrocytes (Fig. 1A) with high penetrance (91.1 ± 2.4%, Fig. 1B) and specificity (96.3 ± 0.9%, Fig. 1C). Penetrance in NeuN positive cells was low (3.6 ± 0.8%, Fig. 1D).

### 3.2. G_i_ pathway activation in ACC astrocytes impairs PA learning

PA learning is hippocampus dependent. Training rats with several PAs leads to schema formation which is stored in the ACC and the learnt PAs gradually become hippocampus independent (Tse et al., 2007; Tse et al., 2011; Hasan et al., 2019; Liu, S. et al., 2022). Astrocytic G_i_ pathway activation was shown to modulate different cognitive functions (Jones et al., 2018; Kol et al., 2020; Liu, S. et al., 2022). However, the effect of ACC astrocytic G_i_ activation on schema memory is yet unknown. To investigate this, bilateral injection of AAV8-GFAP-hM4Di-mCherry into the ACC of rats (n=7) was done to express hM4Di-mCherry in the astrocytes. This group of rats received intraperitoneal (I.P.) CNO (3 mg/kg body weight) 30 mins before the start of each session and 30 mins after the end of each session. Hereafter, this group will be referred as hM4Di-CNO group. Another group of rats (n=6) was used as control which did not receive AAV8-GFAP-hM4Di-mCherry injection and CNO.

After habituation and pretraining, we trained both groups of rats with six PAs (session 1, 2, 4-8, 10-17) (Fig. 2A). Control group showed a gradual increase in performance index (PI) throughout the PA sessions (Fig. 2A). At S6, the mean PI was significantly increased above the chance level (63%, *p*=0.002, one-sample t test with hypothetical value of 50%) and it remained above the chance level throughout the following PA sessions. At S17 the mean PI reached the maximum level (77%). This result is consistent with the other reports (Tse et al., 2007; Hasan et al., 2019; Liu, S. et al., 2022). However, the hM4Di-CNO group consistently had lower mean PI (Fig. 2A) compared to the control group in all PA training sessions from S6-S17 (statistical data is given in the Supplementary Table-3A). At S6, the mean PI of this group was 50% and at S17 it was 58%. This finding indicated that G_i_ pathway activation in ACC astrocytes during and immediately after PA training sessions impaired PAs learning.

### 3.3. G_i_ pathway activation in ACC astrocytes reduces cAMP and L-lactate levels in ACC

Astrocyte derived L-lactate or exogenous L-lactate was shown to confer beneficial effects in cognitive functions in several studies (Newman et al., 2011; Suzuki, Akinobu et al., 2011; Wang, J. et al., 2017; Harris et al., 2019; Vezzoli et al., 2020). As hM4Di is a G_i_-coupled receptor, its activation by CNO could lead to inhibition of adenylyl cyclase resulting in a decreased level of cyclic adenosine monophosphate (cAMP) (Jones et al., 2018). cAMP in astrocytes acts as a trigger for L-lactate production (Choi et al., 2012; Zhou et al., 2021). We hypothesized that hM4Di activation in ACC astrocytes could lead to a decrease in cAMP with a consequent decrease in L-lactate level in ACC. To confirm this, we prepared a cohort of eight rats by habituation and pre-training for PA experiments (Fig. 3A). Then bilateral injection of AAV8-GFAP-hM4Di-mCherry into ACC was done in these rats. After three weeks, all rats were trained for two PA training sessions with six PAs. In S3, rats were given I.P. CNO (3 mg/kg body weight, n=4 rats) or saline (n=4 rats). After 30 mins, PA training was started, and the rats were sacrificed at 60 mins of CNO or saline administration. Brain was collected and immunohistochemistry (IHC) was done to assess the cAMP level. As shown in Fig. 3B and Fig. 3C, cAMP was reduced in the hM4Di expressed cells in CNO treated rats compared to the saline treated rats. Colocalization analysis (Fig. 3C) of hM4Di-mCherry with cAMP showed decreased Mander’s coefficient M1 (ratio of cAMP intensity colocalized with hM4Di-mCherry to total cAMP intensity) in CNO treated rats compared to the saline treated rats (25.1 ± 1.6 vs 32.6 ± 1.9, respectively; t=6.01, df=6, *p*=0.001, unpaired Student’s t test).

Most cells including astrocytes that generate cAMP can export a portion of it into the extracellular fluid (ECF) (Stone & John, 1990; Rosenberg, 1992). Extracellular cAMP levels correlate with the intracellular cAMP levels and could be used to indirectly assess the intracellular cAMP levels and therefore the activity of adenylyl cyclase in the awake, freely moving animals (Stone & John, 1990; Nomikos et al., 2000; Klamer et al., 2005). To detect the effect of G_i_ activation of ACC astrocytes on the ECF cAMP levels at different timepoints, another cohort of eight rats were prepared similarly and were given CNO (n=4) or saline (n=4) I.P. at S3 (Fig. 3A). We collected ECF from ACC by microdialysis before, 20, 40, and 60 minutes after the CNO or saline administration. As shown in Fig. 3D, we observed a significant reduction in cAMP from baseline in the ACC ECF at these timepoints after CNO injection. These results indicated that the reduction in cAMP due to astrocytic G_i_ activation could be observed as early as 20 mins after intraperitoneal administration of CNO in hM4Di expressed rats and the decrease is sustained at least until 60 mins after CNO administration. We also measured the L-lactate levels in these microdialysate samples of ACC. As shown in Fig. 3E, we observed a significant decrease in the L-lactate level in ACC at 20, 40, and 60 minutes after CNO injection compared to the saline suggesting a decreased L-lactate release from astrocytes due to G_i_ activation.

### 3.4. Administration of exogenous L-lactate can rescue the astrocytic G_i_ pathway activation-mediated impairment in PA learning

Given that the G_i_ activation in ACC astrocytes decreases L-lactate levels in ACC, we reasoned that the impaired PA learning observed in the astrocytic G_i_ pathway-activated rats could be rescued by exogenous L-lactate administration if indeed the impairment was due to decreased L-lactate level. To investigate this, we prepared another group of rats (n=7). These rats received bilateral injection of AAV8-GFAP-hM4Di-mCherry into the ACC to express hM4Di-mCherry in the astrocytes. They also received CNO 30 mins before the start and 30 after the end of each PA training session (similar to the hM4Di-CNO group). Moreover, they received exogenous L-lactate bilaterally (10 nmol, 1 μl/ACC) into the ACC 15 mins after receiving CNO injections. Hereafter, this group of rats will be referred as rescue group.

As shown in Fig. 2A, the rescue group showed consistently higher mean PI than the hM4Di-CNO group. Interestingly, their PI was even higher than the control group in S6-S15 (statistical data is given in the Supplementary Table-3B). At S6, the mean PI was 72% and rapidly reached to 78% at S11 and remained ≥78% thereafter suggesting that a smaller number of PA training sessions were required to learn the six PAs in this group compared to the control group. These findings confirmed that the impaired PA learning observed in the hM4Di-CNO group was due to decreased L-lactate level in the ACC upon astrocytic G_i_ pathway activation. It also suggested that exogenous L-lactate not only rescues the astrocytic G_i_ activation-mediated PA learning impairment but can also reduce the number of required PA training sessions to learn the original six PAs.

### 3.5. G_i_ pathway activation in ACC astrocytes impairs memory retrieval whereas concurrent exogenous L-lactate administration rescues the impairment

Non-rewarded probe tests (PT) were performed at S3, S9, and S18 to test the PA memory retrieval. Fig. 2B-D show the results of PT1-3, respectively. In PT1 (Fig. 2B), no rat group spent significantly higher time digging cued sand well above the chance level. In both PT2 (Fig. 2C) and PT3 (Fig. 2D), the control group spent significantly higher time digging cued sand well above the chance level indicating that they learned the PAs from the previous PA training sessions and were able to retrieve it during the non-rewarded PTs (PT2: one sample t test, t=2.95, df=4, *p*=0.042; PT3: one sample t test, t=5.37, df=5, *p*=0.003). Moreover, the digging time percentage in the cued sand well was significantly higher in the PT3 compared to the PT2 indicating further PA learning through S10-S17 after PT2 (percent digging time in cued sand well 21.1 ± 3.1% and 40.3 ± 10.8% in PT2 and PT3 respectively, paired t test, t=4.4, df=4, *p*=0.012). However, the hM4Di-CNO group did not spend higher time digging cued sand well above the chance level in PT2 and PT3. This could be due to the effect of astrocytic G_i_ stimulation itself during the PTs (as these rats received CNO before the PTs) or due to the fact that these rats did not learn the PAs in the first place as evidenced by their poorer performance index during the PA training sessions compared to control rats. To clarify this, both PT2 and PT3 were done twice in the rescue group (Fig. 2C and 2D). One test was done with only CNO in the morning and another test was done with CNO+L-lactate in the afternoon. We found that the rescue group could not retrieve PA memory if only CNO was given. However, they could retrieve PA memory if L-lactate is given concurrently with CNO as evidenced by the significantly higher digging time spent in cued sand well above the chance level (PT2: percent digging time in cued sand well 27.8 ± 6.6, one sample t test, t=4.32, df=6, *p*=0.005; PT3: percent digging time in cued sand well 32.1 ± 5%, one sample t test, t=8.06, df=6, *p*<0.001). This result suggested that G_i_ pathway activation in ACC astrocytes can impair retrieval of already learned PAs and concurrent exogenous L-lactate administration can successfully rescue the impairment in memory retrieval.

### 3.6. G_i_ pathway activation in ACC astrocytes impairs new PA learning despite the existence of prior associative schema whereas exogenous L-lactate administration rescues the impairment

Rats that have prior learning of associative schema showed rapid acquisition of new PAs in a single trial (Tse et al., 2007; Hasan et al., 2019; Liu, S. et al., 2022). We replaced two of the six original PAs (OPAs) with two new PAs (NPAs) at S19 (Fig. 2A) and performed non-rewarded PT4 (PT4) after 24 hours (Fig. 2E). Consistent with other reports (Tse et al., 2007; Hasan et al., 2019; Liu, S. et al., 2022), the control group was able to learn the new PAs from the single PA training session (PI 72.5 ± 3.2%) and retrieve it during the PT4 as evidenced by the significantly higher digging time spent at the correct new cued location (percent digging time in new cued sand well 44.6 ±13.2%, one sample t test, t=5.14, df=5, *p*=0.004). Similarly, the rescue group also learned the new PAs from the single PA training session (PI 70.4 ± 5.5%) and retrieve it during the PT4 as evidenced by the significantly higher digging time spent at the correct new cued location when the PT4 was done with CNO+L-lactate (percent digging time in new cued sand well 36.5 ± 5%, one sample t test, t=10.24, df=6, *p*<0.001). This result confirmed that the rescue group developed the associative schema like control group and can assimilate the new PAs into the existing schema in a single trial if CNO+L-lactate is given during the PA training session (S19). However, PT4 without concurrent L-lactate administration (i.e., with only CNO) showed impaired memory retrieval consistent with the results of PT2 and PT3 of this group further supporting that ACC astrocytic G_i_ activation impairs memory retrieval and exogenous L-lactate administration can rescue the retrieval impairment.

New PA learning requires activation and retrieval of existing associative schema stored in the ACC. We reasoned that G_i_ pathway activation in ACC astrocytes might impair new PA learning even in rats having associative schema memory due to G_i_ pathway activation mediated impairment of memory retrieval from the ACC. To test this hypothesis, we used five rats from the rescue group for further study (Fig. 2F-H). In S21, we checked the mean PI of these rats by original six PAs without giving CNO or L-lactate. The mean PI was 75.3%. In S22, we replaced two old PAs with two new PAs (NPA 9 & 10) and performed PA training with CNO only. The mean PI dropped to 63.3%. Then we performed PT5 with CNO+L-lactate (Fig. 2G). The rats did not spend significantly higher time in digging the new cued sand well than the chance level. This indicated that even though these rats already had associative schema memory, they could not learn the new PAs from single trial due the astrocytic G_i_ pathway activation in ACC during the training session with new PAs.

Next, we examined whether these rats could learn the new PAs if we increase the number of training sessions. With the same PAs as in S22, we continued to do three more PA training sessions (S24-26) with CNO. As shown in the Fig. 2F, the mean PIs in these sessions (69.3%, 66%, and 66% respectively) remained similar to the mean PI of S22 suggesting that the rats were not learning the new PAs despite multiple training sessions. However, at S27, we administered CNO+L-lactate and the mean PI raised to 74%. This training session was followed by PT6 (Fig. 2G) with CNO+L-lactate. The rats spent significantly higher time in digging the new cued sand well above the chance level (percent digging time in new cued sand well 29.9 ± 7.2%, one sample t test, t=4.02, df=4, *p*=0.016) indicating that the rats learned the NPA 9 & 10 from S27. The results suggested that exogenous administration of L-lactate can rescue the impaired new PA learning ability of ACC astrocytic G_i_ pathway activated rats.

Next, we investigated whether these rats can recall OPAs if ACC astrocytic G_i_ pathway is activated, and no exogenous L-lactate is given. In S29 and S30, we checked the mean PI of rats by injecting both CNO and L-lactate (Fig. 2F). Similar to the mean PIs in S6-S17, the mean PIs in these two sessions were 77.33% and 76.67%, respectively. S31-S33 were done with CNO only. In these sessions, the mean PIs dropped to 65.3, 63.3, and 64.7 % respectively indicating poorer performance without exogenous L-lactate which is similar to the mean PIs of the hM4Di-CNO group (S6-S17). In S34, PT7 was done twice; once with only CNO, and once with CNO+L-lactate (Fig. 2H). The rats could not recall the existing associative schema memory if L-lactate was not given in addition to CNO. This result indicated that even after PA learning and schema formation, astrocytic G_i_ pathway activation in ACC can impair PA memory retrieval highlighting the crucial role of ACC astrocyte in schema memory retrieval.

### 3.7. CNO application itself has no effect on PA learning and memory retrieval

Although CNO had long been considered biologically inert, studies showed that it is converted to clozapine. CNO was implicated in reduced startle response to loud acoustic stimuli and clozapine-like interoceptive stimulus effects in rodents (MacLaren et al., 2016; Manvich et al., 2018). Therefore, we investigated whether CNO itself had effect on PA learning, schema formation, and memory retrieval. Rats (n=4) were bilaterally injected with AAV8-GFAP-mCherry into the ACC. After habituation and pretraining, these rats were similarly trained for PA learning. Before 30 mins and after 30 mins of each PA training session, they received I.P. CNO. As shown in Supplementary Fig. S2, the CNO did not affect the PA learning, schema formation, memory retrieval, NPA learning and retrieval, and latency (time needed to commence digging at the correct well). They behaved similar to the control group. This result is consistent with our recent study where CNO did not affect the PA learning and schema formation in rats bilaterally injected with AAV8-GFAP-mCherry into CA1 of hippocampus (Liu, Shu et al., 2022).

### 3.8. G_i_ pathway activation in ACC astrocytes reduces neuronal mitochondrial biogenesis whereas concurrent exogenous L-lactate administration rescues it

Mitochondrial dysfunction is a hallmark of numerous diseases that cause cognitive decline, for example neurodegenerative diseases, genetic mitochondrial diseases, and aging (Golpich et al., 2017; Khacho et al., 2017). Multiple recent studies have provided striking evidence of the role of mitochondrial biogenesis in hippocampal-dependent cognitive functions (Khacho et al., 2017; Liu, Y. et al., 2018; Han et al., 2020; Jacobs et al., 2021). A recent study has demonstrated that exercise-induced L-lactate release from skeletal muscle or I.P. injection of L-lactate can induce hippocampal PGC-1α (Peroxisome proliferator-activated receptor-gamma coactivator 1-alpha) expression and mitochondrial biogenesis in mice (Park et al., 2021). Therefore, we hypothesized that the schema memory impairment observed in our study could be associated with impaired mitochondrial biogenesis caused by the decreased L-lactate levels in ACC upon astrocytic G_i_ activation.

In a proteomic study of hippocampus in our lab, we recently found increased expression of SIRT3 (sirtuin 3) in the hippocampus of anesthetized rats one hour after bilateral administration of exogenous L-lactate into hippocampus compared to ACSF. In Supplementary Fig. S3, western blot of hippocampal protein extract from anesthetized rats one hour after bilateral administration of ACSF or L-lactate (100 nmol of L-lactate dissolved in 1μl of ACSF at a flow rate of 0.1 μl/min in each hippocampus) is given which shows increased expression of SIRT3 in the hippocampus of L-lactate treated rats. SIRT3 is known to promote mitochondrial biogenesis, reduce reactive oxygen species (ROS) production, and plays important role in learning and memory (Fu et al., 2012; Ansari et al., 2017; Satoh et al., 2017; Kim et al., 2019; Liu, Y. et al., 2019; Liu, Q. et al., 2021; Sun et al., 2021). PGC-1α was shown to activate SIRT3 promoter (Sun et al., 2021). On the other hand, SIRT3 was shown to promote PGC-1α expression (Fu et al., 2012) suggesting a positive feedback loop between SIRT3 and PGC-1α. As we have shown that ACC astrocytic G_i_ activation decreases L-lactate in the ACC ECF, we hypothesized that the PGC-1α/SIRT3/mitochondrial biogenesis axis could have been downregulated in the ACC neurons in the hM4Di-CNO group of rats. Fig. 4 shows the results from the control, hM4Di-CNO, and rescue groups of rats used for schema experiments in the current study. The control rats did not receive CNO or L-lactate before being sacrificed. hM4Di-CNO group received I.P. CNO one hour before being sacrificed. Rescue group received I.P. CNO 75 mins before and bilateral exogenous L-lactate into ACC 60 mins before being sacrificed. We observed a significantly decreased expression of PGC-1α, SIRT3, and ATPB (a component of mitochondrial membrane ATP synthase) in the ACC neurons of hM4Di-CNO group compared to the control group. The relative mtDNA copy number in ACC was also decreased in the hM4Di-CNO group (Fig. 4G). On the other hand, the rescue group showed increased expression of PGC-1α, SIRT3, and ATPB in the ACC neurons as well as increased relative mtDNA copy number in ACC even higher than the control group which is consistent with their higher performance in the PA training than the control group. Together, these results revealed that ACC astrocytic G_i_ activation impairs neuronal mitochondrial biogenesis by decreasing ECF L-lactate levels in ACC and that exogenous L-lactate administration rescues the impaired mitochondrial biogenesis.

**Fig. 4.**
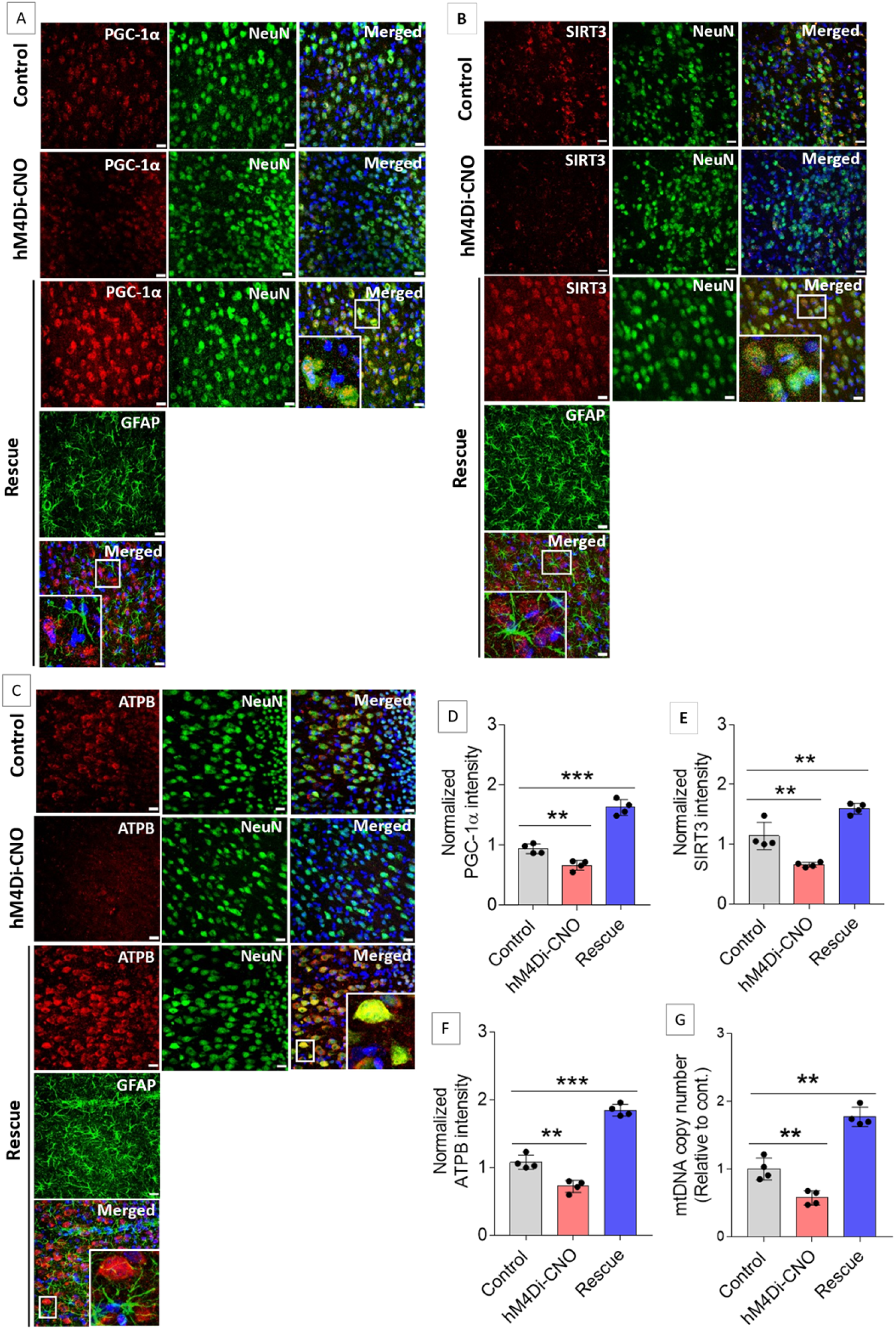
G_i_ activation in ACC astrocytes reduces neuronal mitochondrial biogenesis whereas concurrent exogenous L-lactate administration rescues the impairment. **A-C.** Representative confocal micrograph of PGC-1α (**A**)/ SIRT3 (**B**)/ ATPB (**C**) co-labelled with NeuN, GFAP, and DAPI in the ACC of control, hM4Di-CNO, and rescue groups from schema experiments. Astrocytic G_i_ pathway activation (hM4Di-CNO group) in ACC resulted in decreased of PGC-1α/ SIRT3/ ATPB expression in ACC, whereas concurrent exogenous L-lactate (rescue group) administration resulted in increased PGC-1α/ SIRT3/ ATPB expression. Scale bars: 20 μm. **D-F.** Fluorescence intensity of PGC-1α (**D**)/ SIRT3 (**E**)/ ATPB (**F**) stained sections in the ACC of hM4Di-CNO and rescue groups were assessed and normalized to control group of rats. Data is shown as mean ± SD (n=4 rats per group). \*\*\**p*<0.001, \*\**p*<0.01, unpaired Student’s t-test. **G.** mtDNA copy number abundance in the ACC of control, hM4Di-CNO and rescue groups relative to nDNA. Relative mtDNA copy number was significantly decreased in hM4Di-CNO group, whereas it was increased in the rescue group compared to control. Data are shown as mean ± SD (n=4 rats per group). \*\**p*<0.01, unpaired Student’s t-test.

### 3.9. Mitochondrial biogenesis by L-lactate is dependent on MCT2 and NMDAR activity

Previous studies demonstrated that L-lactate entry into the neurons is needed for its beneficial effect (Newman et al., 2011; Suzuki, A. et al., 2011; Wang, J. et al., 2017). After entry, L-lactate promotes plasticity gene expression by potentiating NMDA signaling (Yang et al., 2014; Magistretti & Allaman, 2018). To investigate whether entry into the neuron and NMDAR activity is required for L-lactate induced mitochondrial biogenesis, we prepared 20 rats as shown in the Fig. 5A. After habituation and pretraining, cannula placement was done bilaterally into the ACC. After one week of recovery, two PA training sessions (S1 and S2) with six OPAs were done. At S3, rats were randomly placed into one of the five groups (n=4 rats in each group): ACSF, L-Lactate, 4-CIN+L-Lactate, D-APV, D-APV+L-Lactate. ACSF group received bilateral infusion of ACSF into ACC (1 μl), then placed into the PA event arena for 60 mins after which rats were sacrificed and ACC were collected for relative mtDNA copy number analysis. Similarly, L-Lactate and D-APV groups received bilateral infusion of L-lactate (10 nmol, 1 μl) or D-APV (30 mM, 0.5 μl) into the ACC and sacrificed after 60 mins. D-APV is a competitive inhibitor of glutamate binding site of NMDA receptor. 4-CIN+L-Lactate group received bilateral infusion of α-Cyano-4-hydroxycinnamic acid (4-CIN, an MCT2 blocker) (0.5 nmol, 1 μl) into the ACC to block neuronal MCT2 to prevent L-lactate’s entry into the neurons. After 15 mins, bilateral L-lactate (10 nmol, 1 μl) infusion into ACC was done and rats were sacrificed after 60 mins. D-APV+L-Lactate group received bilateral infusion of D-APV (30 mM, 0.5 μl) into the ACC to inhibit NMDA receptor. After 15 mins, bilateral L-lactate (10 nmol, 1 μl) infusion into ACC was done and rats were sacrificed after 60 mins.

**Fig. 5.**
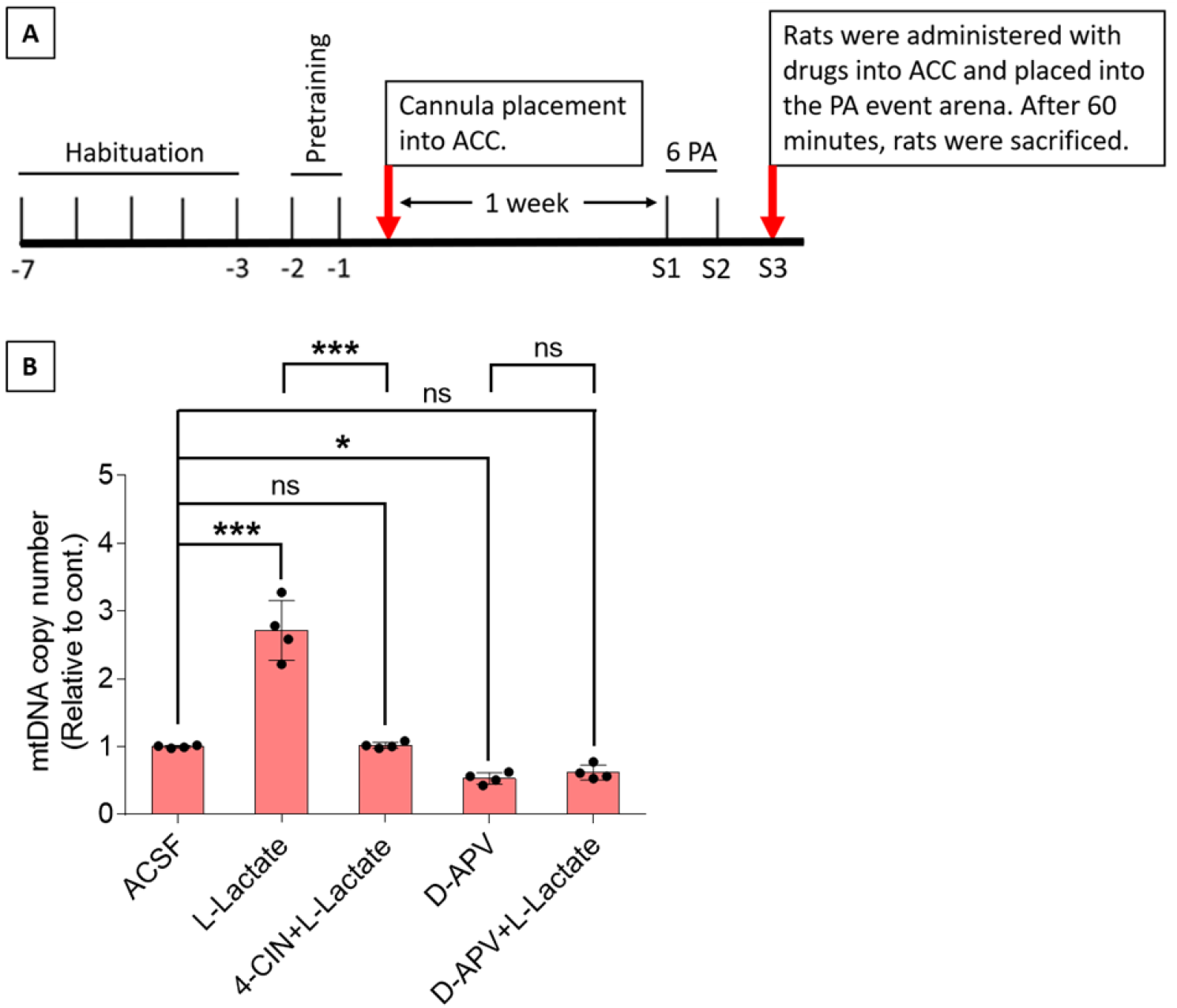
Mitochondrial biogenesis by L-lactate is dependent on MCT2 and NMDAR activity. **A.** Experimental design to investigate whether entry into the neuron and NMDAR activity is required for L-lactate induced mitochondrial biogenesis. **B.** mtDNA copy number abundance in the ACC of ACSF, L-Lactate, 4-CIN+L-Lactate, D-APV, D-APV+L-Lactate groups relative to nDNA. Data are shown as mean ± SD (n=4 in each group). \*\*\**p*<0.001, \**p*<0.05, ns=not significant, ANOVA followed by Tukey’s multiple comparisons test.

As shown in Fig. 5B, L-lactate group showed significantly increased relative mtDNA copy number compared to the ACSF group (*p*<0.001, ANOVA followed by Tukey’s multiple comparisons test). However, this effect was completely abolished if entry into the neurons is blocked by 4-CIN (4-CIN+L-lactate group). Furthermore, inhibition of NMDAR by D-APV (D-APV group) decreased relative mtDNA copy number compared to the ACSF group which cannot be rescued by exogenous L-lactate administration (D-APV+L-lactate group). Together, these results uncovered that L-lactate’s entry into the neuron and NMDAR activity is required for L-lactate mediated mitochondrial biogenesis - revealing another NMDAR dependent signaling effect of L-lactate besides plasticity gene expression.

## 4. Discussion

There has been a paradigm shift in neuroscience in which animal behavior is now considered as a result arising from the coordinated activity of neurons and glia especially astrocytes rather than a result exclusively from neuronal activity (Kofuji & Araque, 2021a). A core feature of astrocytes across CNS is the expression of GPCR genes (Endo et al., 2022), however, astrocytic responses to GPCR activation may show heterogeneity among brain regions or even within a brain region (Kofuji & Araque, 2021b). GPCRs confer astrocytes with the ability to sense synaptic activity and respond with gliotransmitters to regulate neuronal and synaptic functions (Kofuji & Araque, 2021b). L-lactate, derived primarily from astrocytes in the brain, has increasingly been recognized as a novel gliotransmitter that facilitates neuronal plasticity and cognitive functions (Newman et al., 2011; Suzuki, Akinobu et al., 2011; Wang, J. et al., 2017; Magistretti & Allaman, 2018; Harris et al., 2019; Vezzoli et al., 2020). Our current study shows that astrocytic G_i_ activation in ACC decreases intracellular cAMP and ECF L-lactate levels in the ACC.

Schema is defined as a framework of knowledge. New learning occurs rapidly if it occurs against a background of established relevant schema. Rats trained with multiple flavor-place PAs develop schema that enables rapid assimilation of new PA learning (Tse et al., 2007; Tse et al., 2011; Hasan et al., 2019; Liu, S. et al., 2022). Previous studies suggested that PA learning is hippocampus dependent and the associative schema is stored in the ACC (Tse et al., 2007; Tse et al., 2011). We previously demonstrated that neuronal inhibition in either hippocampus or ACC impaired PA learning and schema formation (Hasan et al., 2019). The study also showed that ACC myelination is necessary for PA learning and schema formation and that repeated PA learning is associated with oligodendrogenesis in ACC. However, role of astrocytes in this context was unknown. The current study has showed that ACC astrocytic G_i_ activation impaired PA learning and schema formation, PA memory retrieval, and new PA learning by decreasing L-lactate level in ACC. These impairments were rescued by exogenous L-lactate administration directly into the ACC. Taken together with our previous study (Hasan et al., 2019), our results contribute to the mounting evidence of the glial contribution in cognitive functions and underscores the new paradigm in which glial cells are not considered merely supporting cells to neurons rather a crucial player in cognitive functions along with the neurons. Disruption of neurons, or myelin which is formed by oligodendrocytes, or astrocytes in ACC can disrupt schema memory consolidation. Furthermore, in another study (Liu, S. et al., 2022), we have recently demonstrated that G_i_ activation in the CA1 astrocytes impairs PA learning and schema formation by disrupting recruitment of CA1-ACC projecting neurons highlighting the importance of normal functioning of hippocampal astrocytes in PA learning and schema formation. These discoveries have clinical implications as they suggest that pathological processes involving any of these cell types can eventually result in loss of harmony among these cells and manifest as cognitive impairments. Indeed, accumulating evidence suggests the crucial contributions of non-neuronal cells in the pathology of diverse neurodegenerative disorders including Alzheimer disease, Parkinson disease, Huntington disease, and amyotrophic lateral sclerosis (Brandebura et al., 2022).

In the current study, we have demonstrated that ACC astrocytic G_i_ activation impairs new PA learning even if a prior associative schema exists. This impairment is the result of impaired activation and retrieval of prior associative schema from ACC neuronal network mediated by decreased L-lactate in ACC as exogenous L-lactate administration abolished the impairment. After an associative schema is formed in ACC due to repeated PA training with multiple PAs, the effect of astrocytic G_i_ activation in ACC during new PA learning is different from the effect of astrocytic G_i_ activation in the hippocampus. Whereas G_i_ activation in ACC leads to impairment in both PA memory retrieval and new PA learning as shown in the current study, G_i_ activation in hippocampus primarily affects the new PA learning but not the memory retrieval of the previously learned PAs (Liu, S. et al., 2022). This indicates that once associative schema is formed in the ACC, it becomes independent of hippocampus, and disruption of either hippocampal neuronal (Hasan et al., 2019) or astrocytic (Liu, S. et al., 2022) functions does not impact retrieval of the previously learned PAs. However, new PAs learning in the setting of existence of prior associative schema can be impaired by disruption of either hippocampal functions [neuronal inhibition (Hasan et al., 2019) or astrocytic G_i_ activation (Liu, S. et al., 2022)] or ACC functions [neuronal inhibition (Hasan et al., 2019) or astrocytic G_i_ activation (current study)] indicating that new PA learning requires simultaneous activation of both hippocampus and ACC with optimal functioning of both neurons and astrocytes.

L-lactate is used both as an energy substrate and a signaling molecule. After entry into neuron, it can be converted to pyruvate by lactate dehydrogenase (LDH) during which NADH is also produced. Pyruvate can enter the mitochondria and be processed through Kreb’s cycle and oxidative phosphorylation to generate ATP. Demand of ATP is high in active neurons to maintain various physiological activities including neural plasticity and memory formation (Magistretti & Allaman, 2018). Although astrocyte-derived or exogenous L-lactate’s beneficial effect in cognitive functions have been clearly demonstrated in the current study and several other previous studies (Newman et al., 2011; Suzuki, Akinobu et al., 2011; Wang, J. et al., 2017; Harris et al., 2019; Vezzoli et al., 2020), whether the beneficial effect is conferred by its usefulness as an energy substrate has been debated (Diaz-Garcia et al., 2017). On the other hand, L-lactate’s signaling role is being increasingly recognized. NADH produced during conversion of L-lactate into pyruvate can induce plasticity gene expression by activating NMDA and/or MAPK signaling pathway (Yang et al., 2014; Magistretti & Allaman, 2018; Margineanu et al., 2018). In the current study, we showed that L-lactate induces mitochondrial biogenesis which is dependent on MCT2 and NMDAR activity – uncovering yet another NMDAR dependent signaling role of L-lactate besides plasticity gene expression (Fig. 6). The current study demonstrates that ACC astrocytic G_i_ activation impairs PAs learning, schema formation, and memory retrieval that were associated with a downregulation of neuronal PGC-1α, SIRT3, and mitochondrial biogenesis in ACC. SIRT3, a member of sirtuin family, is a protein deacetylase that is exclusively found in mitochondria and is known to promote mitochondrial biogenesis and reduce ROS production (Fu et al., 2012; Ansari et al., 2017; Sun et al., 2021). Several recent studies have demonstrated that SIRT3 plays important role in learning and memory (Kim et al., 2019; Liu, Y. et al., 2019; Liu, Q. et al., 2021). *Sirt3*^-/-^ mice demonstrated impaired remote memory function and decreased synaptic plasticity and neuronal number in the ACC (Kim et al., 2019). Another study in aged mice showed that SIRT3 overexpression can provide protection against anesthesia/surgery-induced synaptic plasticity dysfunction in hippocampus and attenuate hippocampus-dependent cognitive decline (Liu, Q. et al., 2021). SIRT3 was shown to be required for the anxiolytic and cognition-enhancing effects of intermitted fasting (Liu, Y. et al., 2019). The study demonstrated that mice lacking SIRT3 in the hippocampal neurons have heightened anxiety, poor memory retention, and impaired long-term potentiation at hippocampal synapses. Recently, it has been demonstrated that PGC-1α overexpression in neurons can improve hippocampal neuronal function, increase ATP production, reduce oxidative stress, and attenuate cognitive impairment after chronic cerebral hypoperfusion in mice (Han et al., 2020). Another study has suggested that upregulation of PGC-1α and mitochondrial biogenesis in hippocampus enhances spatial learning and short-term memory (Jacobs et al., 2021). Mitochondrial dysfunction was shown to impair hippocampal-dependent learning and memory in mice (Khacho et al., 2017). On the other hand, another study showed that ameliorating mitochondrial dysfunction rescues carbon ion-induced hippocampal cognitive deficits (Liu, Y. et al., 2018). Mitochondrial dysfunction is a hallmark of numerous diseases that causes cognitive decline (Golpich et al., 2017). Collectively, these reports provided striking evidence of the role of mitochondrial homeostasis in cognitive functions. Therefore, the associations observed from the current study set up an interesting premise for further studies to investigate whether L-lactate regulated neuronal mitochondrial biogenesis plays causal role in the cognitive impairment due to astrocytic G_i_ activation. Based on the known functions, one might hypothesize that L-lactate induced upregulation of PGC-1α/SIRT3/mitochondrial biogenesis could enable the neurons to generate more ATP while reducing oxidative stress during bioenergetic challenges as in cognitively demanding tasks of PAs learning. In line with this, recent study has demonstrated that L-lactate causes a mild ROS burst that induces antioxidant defenses and pro-survival pathways (Tauffenberger et al., 2019).

**Fig. 6.**
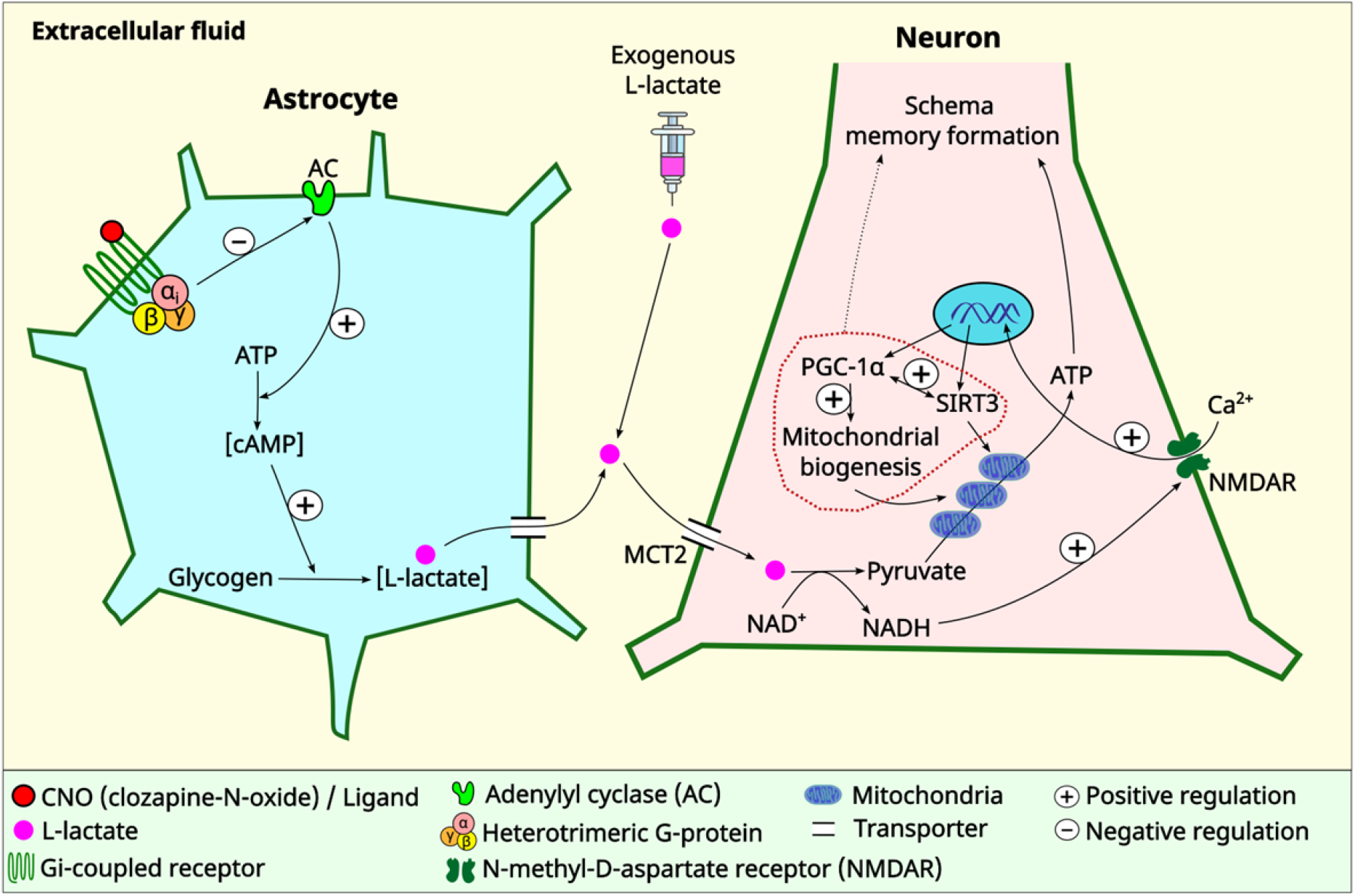
Schematic diagram showing astrocytic G_i_ signaling and L-lactate modulating schema memory and mitochondrial biogenesis. Astrocyte-derived L-lactate in ACC is required for schema memory formation and NMDAR-dependent neuronal mitochondrial biogenesis. Astrocytic G_i_ activation results in decreased L-lactate production with consequent impairments in schema memory and neuronal mitochondrial biogenesis which could be rescued by exogenous L-lactate administration directly into the ACC. Further research is needed to establish the mechanism and the extent of the contribution of mitochondrial biogenesis in schema memory formation (dotted arrow). MCT2: Monocarboxylate transporter 2

In summary, the present study illustrates that ACC astrocytic G_i_ pathway activation impairs schema memory in rats by decreasing L-lactate levels in ACC which was associated with impaired mitochondrial biogenesis in neurons. Furthermore, we demonstrated that L-lactate mediated neuronal mitochondrial biogenesis is dependent on NMDA receptor activity – uncovering a novel signaling mechanism of L-lactate in the brain. Together, these results might have implications in understanding how perturbation in astrocytic functions could impair cognitive functions as well as providing the potential therapeutic targets for ameliorating such impairments.

## Abbreviations

AAV8: Adeno-associated viral vectors serotype 8
ACC: Anterior cingulate cortex
ACSF: Artificial cerebrospinal fluid
ANLS: Astrocyte-neuron L-lactate shuttle
ATP: Adenosine triphosphate
cAMP: Cyclic adenosine monophosphate
CNO: Clozapine N-oxide
DREADD: Designer receptors exclusively activated by designer drug
ECF: Extracellular fluid
GFAP: Glial fibrillary acidic protein
GPCR: G-protein-coupled receptor
MCT: Monocarboxylate transporter
mtDNA: Mitochondrial DNA
nDNA: Nuclear DNA
NeuN: Neuronal nuclei
NPA: New paired associate
OPA: Original paired associate
PA: Paired associate
PBS: Phosphate buffered saline
PFA: Paraformaldehyde
PGC-1α: Peroxisome proliferator-activated receptorgamma coactivator 1-alpha
PI: Performance index
PT: Probe test
ROS: Reactive oxygen species
SIRT3: Sirtuin 3

## CRediT authorship contribution statement

Mastura Akter: Conceptualization; Methodology; Investigation (Schema, stereotactic surgical procedures, microdialysis, IHC, WB, confocal imaging, RT-PCR); Formal analysis; Visualization; Writing - Original Draft

Mahadi Hasan: Investigation (Schema, stereotactic surgical procedures, microdialysis)

Aruna Surendran Ramkrishnan: Investigation (Schema)

Xianlin Zheng: Investigation (Stereotactic surgical procedures; IHC)

Zafar Iqbal: Investigation (Stereotactic surgical procedures)

Zhongqi Fu: Investigation (Schema, microdialysis)

Zhuogui Lei: Investigation (Stereotactic surgical procedures)

Anwarul Karim: Methodology; Formal analysis; Visualization

Ying Li: Conceptualization; Methodology; Resources; Writing - Review & Editing; Supervision; Project administration; Funding acquisition

All the authors approved the final version of the manuscript.

## Additional files

### Supplementary material

Contains supplementary figures and tables

### Source data files

Zip file containing the source data of the figures

## Supplementary material

**Supplementary Fig. S1:**
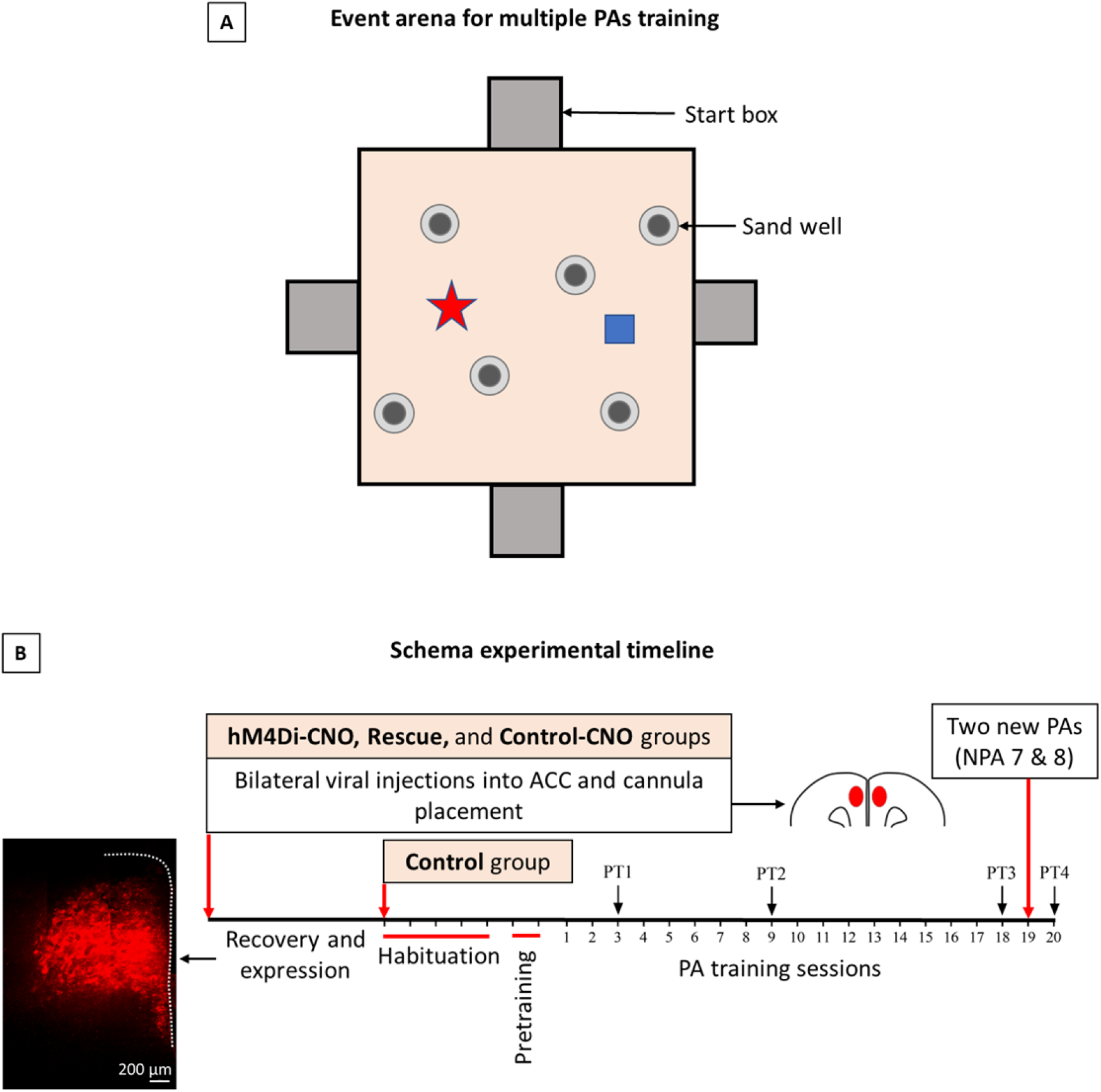
**A**. Event arena for multiple flavor-place paired associate training. **B**. Timeline for schema experiments.

**Supplementary Fig. S2:**
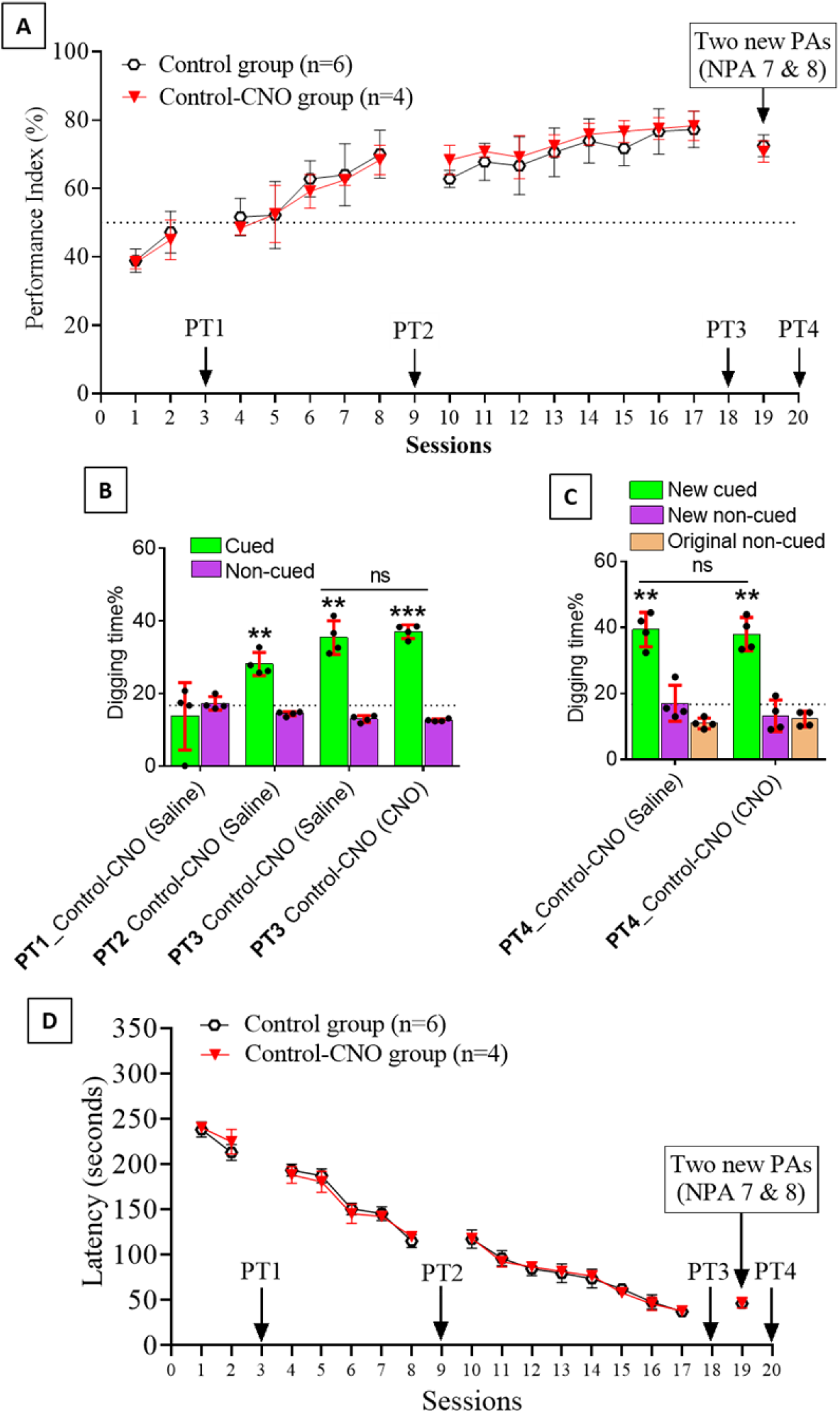
CNO application itself has no effect on PA learning and memory retrieval. **A.** PI (mean ± SD) during the acquisition of the original six PAs (OPAs) (S1-2, 4-8, 10-17) and new PAs (NPAs) (S19) of the control (n=6) and control-CNO (n=4) groups. The control group data is the same data as shown in the Fig. 2. **B.** Non-rewarded PTs (PT1, PT2, and PT3 done on S3, S9, and S18, respectively) to test memory retrieval of OPAs for the control-CNO group. **C.** Non-rewarded PT4 (S20) which was done after replacing two OPAs with two NPAs (NPA 7 & 8) in S19 for the control-CNO group. **D.** Latency (in seconds) before commencing digging at the correct well for control and control-CNO groups. Data shown as mean ± SD. The control group data is the same data as shown in the Fig. 2.

**Supplementary Fig. S3:**
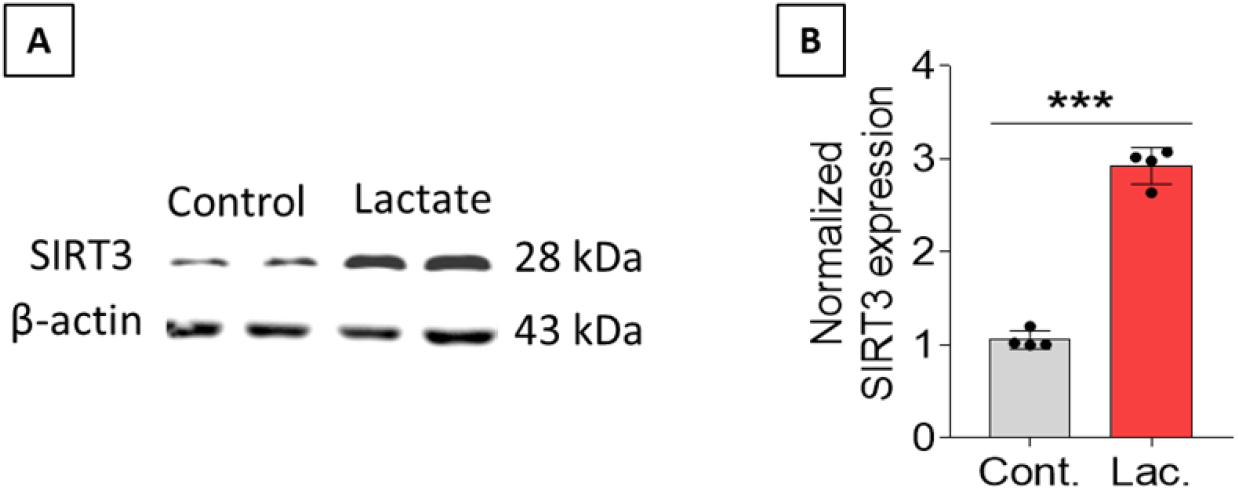
Exogenous L-lactate administration into the hippocampus increases SIRT3 expression. **A.** Representative WB images of SIRT3 expression in the hippocampal protein extracts. Rats were anesthetized and received L-lactate administration bilaterally into the hippocampus (100 nmol of L-lactate dissolved in 1μl of PBS at a flow rate of 0.1 μl/min in each hippocampus) or same volume of ACSF (control). Rats were sacrificed after one hour of L-lactate/ACSF administration and brain was collected for western blot analysis of hippocampus. SIRT3 was found to be significantly increased in the hippocampus of L-lactate treated rats compared to control. **B.** Intensity was quantified and normalized with β-actin. Data are shown as mean ± SD (n=4 rats per group). p***<0.001, unpaired Student’s t-test.

**Supplementary Table-1.** Antibodies used in IHC and WB

| Name of antibodies | Source | Manufacturer and Catalogue number | Dilution |
| --- | --- | --- | --- |
| Anti-cAMP | Rabbit | Abcam_134901 | 1:500 (IHC) |
| Anti-mCherry | Chicken | Abcam_205402 | 1:1000 (IHC) |
| Anti-SIRT3 | Rabbit | SAB5700222 | 1:500 (WB), 1: 250 (IHC) |
| Anti-PGC-1α | Rabbit | Abcam_191838 | 1: 500 (IHC) |
| Anti-ATPB | Mouse | Abcam_14730 | 1:500 (IHC) |
| Anti-β-actin | Mouse | Immunoway_YM3028 | 1:5000 (WB) |
| Anti-GFAP | Mouse | Abcam_4648 | 1:500 (IHC) |
| Anti-NeuN | Rabbit | EMD Millipore corp. USA, AB978 | 1:500 (IHC) |
| Alexa Flour 488, 594 goat anti mouse IgG (H+L) | Goat | Thermo fisher scientific_A11001, Thermo fisher scientific_A11032 | 1:300 (IHC) |
| Alexa Flour 488, 594 goat anti rabbit IgG (H+L) | Goat | Thermo fisher scientific_A11034, Thermo fisher scientific_A11037 | 1:300 (IHC) |
| Goat anti-rabbit secondary antibody, HRP | Rabbit | Invitrogen_31460 | 1:5000 (WB) |

**Supplementary Table-2.**
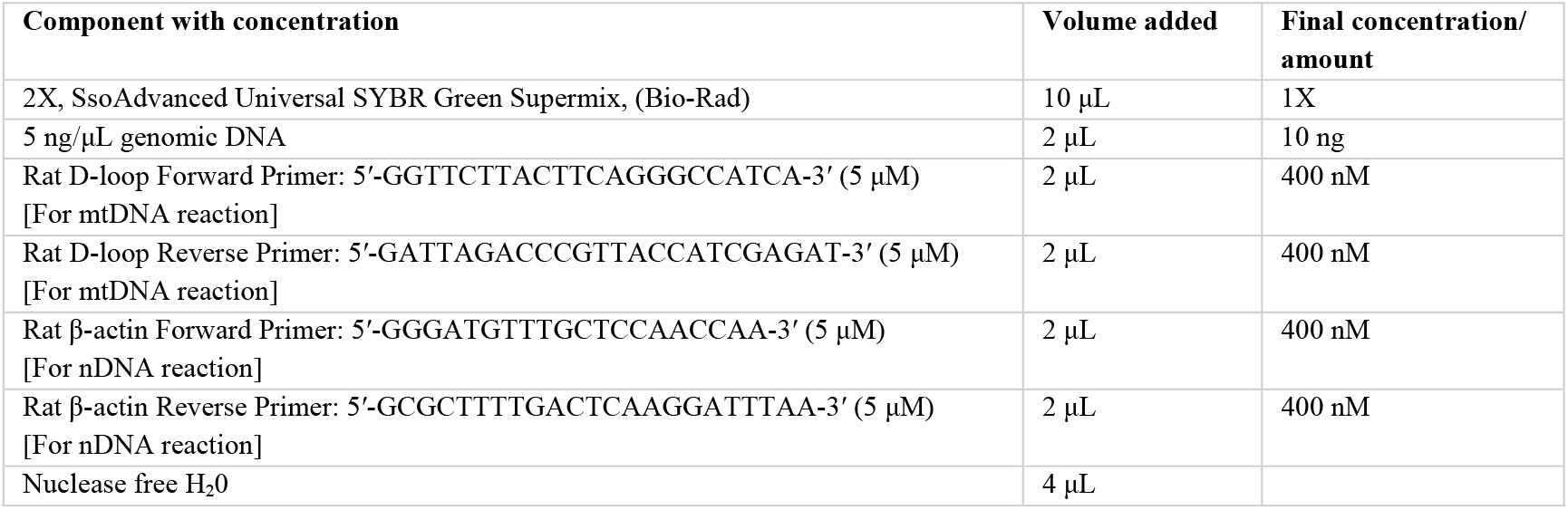
Primer sequences and preparation of 20 μL reaction mixture for real-time PCR

**Supplementary Table-3A.**
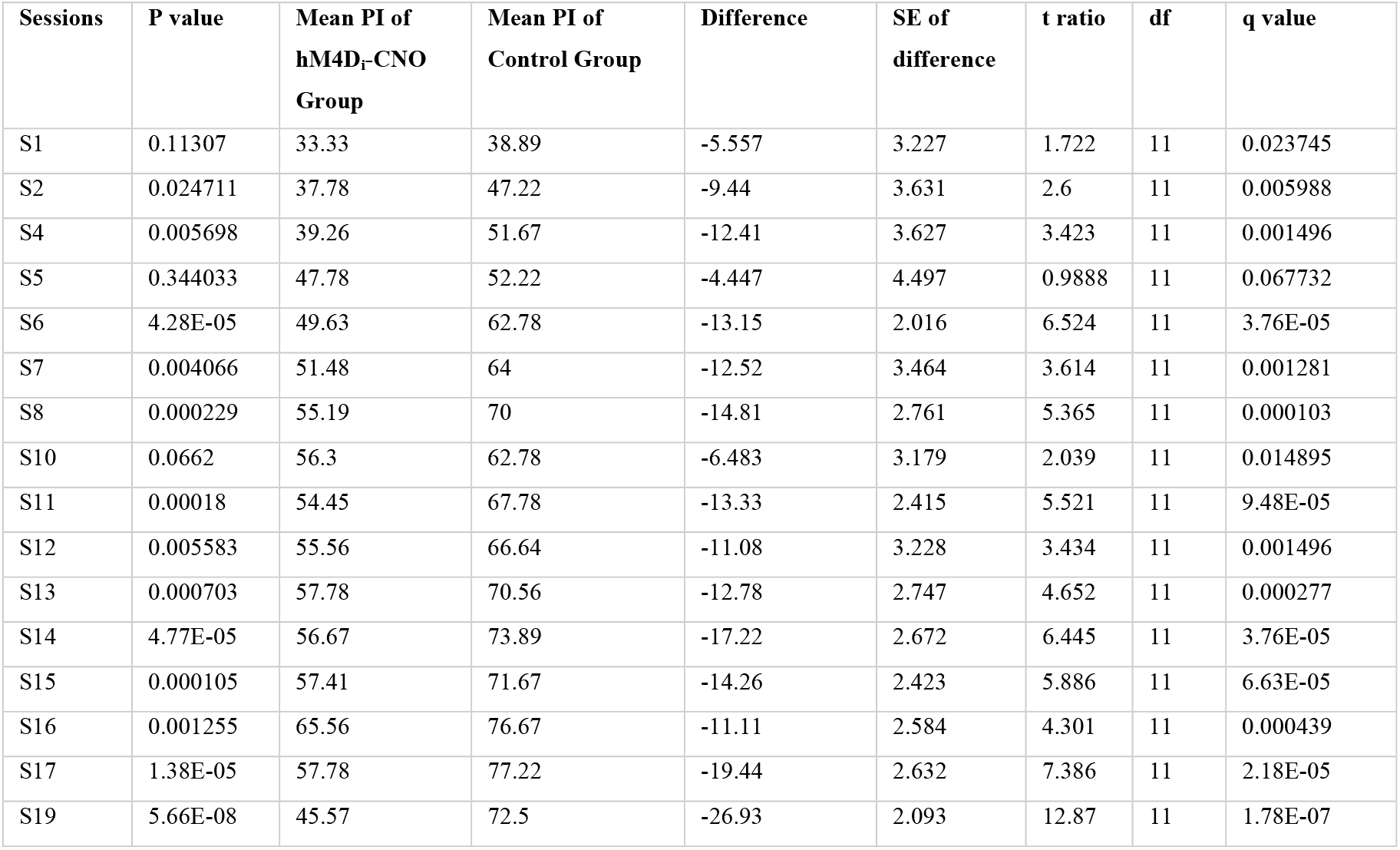
Comparison of performance index of control vs. hM4D_i_-CNO group (Unpaired t test, FDR correction with two-stage step-up method of Benjamini, Krieger and Yekutieli)

**Supplementary Table-3B.** Comparison of performance index of control vs. rescue group (Unpaired t test, FDR correction with two-stage step-up method of Benjamini, Krieger and Yekutieli)

| Sessions | P value | Mean PI of rescue Group | Mean PI of Control Group | Difference | SE of difference | t ratio | df | q value |
| --- | --- | --- | --- | --- | --- | --- | --- | --- |
| S1 | 0.550501 | 40.67 | 38.89 | 1.777 | 2.885 | 0.6159 | 11 | 0.371588 |
| S2 | 0.07656 | 42.33 | 47.22 | -4.887 | 2.501 | 1.954 | 11 | 0.072349 |
| S4 | 0.288362 | 55.56 | 51.67 | 3.887 | 3.484 | 1.116 | 11 | 0.227085 |
| S5 | 0.103651 | 59.11 | 52.22 | 6.887 | 3.881 | 1.774 | 11 | 0.089046 |
| S6 | 0.004271 | 72.22 | 62.78 | 9.443 | 2.633 | 3.586 | 11 | 0.01009 |
| S7 | 0.010441 | 75.33 | 64 | 11.33 | 3.678 | 3.082 | 11 | 0.016445 |
| S8 | 0.030339 | 77.78 | 70 | 7.777 | 3.13 | 2.484 | 11 | 0.031856 |
| S10 | 1.35E-05 | 73.78 | 62.78 | 11 | 1.485 | 7.408 | 11 | 0.000127 |
| S11 | 0.000544 | 78 | 67.78 | 10.22 | 2.125 | 4.811 | 11 | 0.002571 |
| S12 | 0.003173 | 79.55 | 66.64 | 12.91 | 3.437 | 3.757 | 11 | 0.009995 |
| S13 | 0.007587 | 80 | 70.56 | 9.44 | 2.895 | 3.261 | 11 | 0.014339 |
| S14 | 0.018412 | 80.89 | 73.89 | 6.997 | 2.531 | 2.764 | 11 | 0.024856 |
| S15 | 0.025361 | 78.22 | 71.67 | 6.55 | 2.534 | 2.585 | 11 | 0.029958 |
| S16 | 0.668788 | 78 | 76.67 | 1.327 | 3.018 | 0.4395 | 11 | 0.421337 |
| S17 | 0.7724 | 78 | 77.22 | 0.7767 | 2.62 | 0.2965 | 11 | 0.456199 |
| S19 | 0.438051 | 70.44 | 72.5 | -2.057 | 2.556 | 0.8047 | 11 | 0.318429 |

